# DNA-measuring Wadjet SMC ATPases restrict smaller circular plasmids by DNA cleavage

**DOI:** 10.1101/2022.10.04.510915

**Authors:** Hon Wing Liu, Florian Roisné-Hamelin, Bertrand Beckert, Alexander Myasnikov, Yan Li, Stephan Gruber

## Abstract

Structural maintenance of chromosomes (SMC) complexes fold DNA by loop extrusion to support chromosome segregation, genome maintenance, and gene expression. Wadjet systems (JetABCD/MksBEFG/EptABCD) are derivative SMC complexes with proposed roles in bacterial immunity against selfish DNA elements. Here, we show that JetABCD systems restrict extrachromosomal circular DNA with an upper size limit of about 100 kb, while a linear plasmid evades restriction. Recombinant preparations of a JetABCD complex cleave circular DNA regardless of its helical topology but not linear DNA; cleavage occurs at random positions and depends on ATP as well as the SMC ATPase. We solve a structure of the JetABCD core by cryo-EM revealing an alternative dimer-of-dimers configuration. The two SMC DNA motor units face in opposite orientations—rather than the same as observed with MukBEF—possibly representing a restriction-specific state of JetABCD. We hypothesize that JetABCD is a DNA-shape-specific endonuclease and present a model for the activation of DNA cleavage exclusively when extrusion of an entire plasmid has been completed by a single JetABCD complex. The self-stalling of two opposing DNA motor units may serve as signal to trigger DNA cleavage. Complete extrusion by a single complex cannot be achieved on the much larger chromosome, explaining how self-DNA may evade cleavage.

## Introduction

Bacteria rely on horizontal gene transfer (HGT) by DNA conjugation, natural transformation, and phage transduction to quickly adapt to rapidly changing environments. Selfish genetic elements utilize these pathways to spread from cell to cell, often with undesired consequences on fitness, for example due to metabolic burden. Bacteria have evolved an armada of defense systems that counteract selfish elements; those systems that target DNA must be able to differentiate potentially detrimental nonself-DNA from self-DNA to spare the host chromosome(s). CRISPR/Cas systems target specific DNA base sequences that are absent from the chromosome, while restriction-modification systems discriminate on the basis of DNA modifications. More recently, SMC and SMC-like complexes have also been implicated in bacterial immunity against plasmids (Doron et al., 2018; Jaskolska et al., 2022; Panas et al., 2014). A putatively related function in viral restriction against HBV, EBV, KSHV, and HIV-1 has been reported for the Smc5/6 complex in human cells (Decorsiere et al., 2016; Dupont et al., 2021; Han et al., 2022; Murphy et al., 2016; Yiu et al., 2022).

SMC complexes are multi-subunit DNA motor complexes that normally fold chromosomal DNA into a loop or a series of DNA loops in a process called DNA loop extrusion (Yatskevich et al., 2019). They are found in all domains of life. In eukaryotes, three types of complexes (denoted as Smc5/6, cohesin and condensin) are nearly ubiquitous (Yoshinaga and Inagaki, 2021). In prokaryotes, the Smc-ScpAB complex is predominant; it contributes to chromosome organization and segregation in diverse bacteria and likely also in archaea (Gruber, 2018). Other SMC variants, named MukBEF and MksBEF (for MukBEF-like SMC complexes), also support chromosome organization in bacteria (Lioy et al., 2020; Makela and Sherratt, 2020; Petrushenko et al., 2011). MukBEF is found in *E. coli* and related γ-proteobacteria, while MksBEF is widely scattered over the phylogenetic tree, likely as a result of HGT events (Gruber, 2011; Petrushenko *et al*., 2011). A high divergence of Muk and Mks protein sequences from other SMC sequences indicates elevated rates of sequence evolution.

Some MksBEF systems harbor an additional MksG subunit that shows sequence similarity to topoisomerase/primase (TOPRIM) domains (Petrushenko *et al*., 2011). A *Mycobacterium smegmatis* strain harboring a sporadic mutation in the SMC subunit of such a system had unknowingly been selected as lab strain owing to its efficient plasmid transformation, thus implicating MksBEFG in plasmid restriction (Panas *et al*., 2014). Four-membered MksBEFG systems, designated as Wadjet or JetABCD, have since been uncovered in multiple species and shown to restrict plasmids in *B. subtilis* and *Corynebacterium glutamicum* (Bohm et al., 2020; Doron *et al*., 2018). All systems (hereafter denoted as the Wadjet family of JetABCD systems) share the characteristic domain organization with other SMC complexes, suggesting that they support a similar biochemical activity and raising the question of how DNA loop extrusion or DNA translocation may be related to the discrimination of plasmid and chromosome DNA.

Wadjet systems comprise four subunits: the ‘kleisin’ JetA (MksF), the ‘kite’ JetB (MksE), the SMC JetC (MksB), and Wadjet-specific JetD (MksG) (Figure 1A). All four subunits appear essential for plasmid restriction (Doron *et al*., 2018). SMC subunits harbor a long intramolecular coiled coil ‘arm’ with a dimerization ‘hinge’ domain at one end and an ATP Binding Cassette (ABC) ATPase ‘head’ domain at the other (Yatskevich *et al*., 2019). Kite subunits associate as a homodimer with the middle segment of a kleisin subunit. Kleisin proteins typically connect the head of one SMC protein to the head-proximal coiled coil of another, thus forming the typical tripartite protein ring topology (Burmann et al., 2013). MukF-type kleisins moreover undergo homotypic interactions via additional N-terminal sequences, thus generating dimer-of-dimers, with the two associating motor units presumably translocating in opposite direction to support two-sided symmetric DNA loop extrusion (Burmann et al., 2021; Fennell-Fezzie et al., 2005). Like MukF, JetA includes additional amino-terminal sequences implying that the JetABCD complex would also form a dimer-of-dimers (‘d-o-d’).

**Figure 1.**
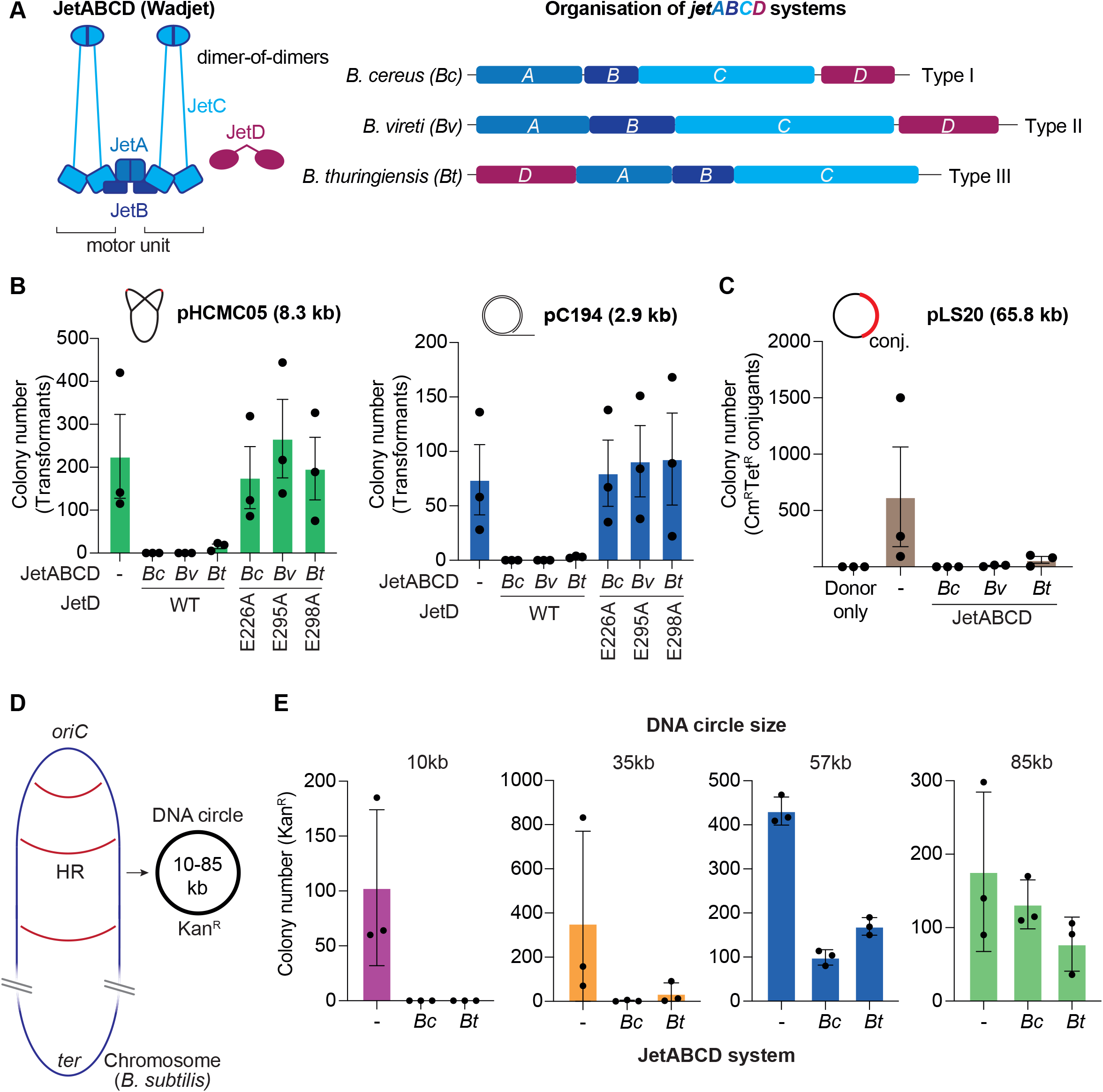
Determinants for plasmid restriction by Bacillus JetABCD *in vivo*. **A)** Left panel: Architecture of a dimer-of-dimers (‘d-o-d’) of JetABCD, with each JetC dimer comprising a motor unit. Right panel: Operon organization for the studied *Bacillus* JetABCD systems. **B)** Plasmid transformations. Counts of antibiotic-resistant colonies for the indicated *B. subtilis* backgrounds after transformation with the indicated plasmids. *Bc*: *B. cereus*; *Bv*: *B. vireti*; *Bt*: *B. thuringiensis*. Means and standard deviations from three independent experiments are shown. **C)** Plasmid conjugation. Counts of chloramphenicol-, tetracycline-resistant transconjugants for the denoted *B. subtilis* recipients (TetR) after conjugation from a donor (CmR) with pLS20(cat). Means and standard deviations from three independent experiments are shown. **D)** Concept of chromosomal excision. **E)** Chromosome excision. Counts of kanamycin-resistant (containing DNA circles) denoted *B. subtilis* strains. Means and standard deviations from three independent experiments are shown. See Figure S1 for further experiments and a more detailed scheme for the chromosome excision assay. See also Figure S1.

DNA loop extrusion by SMC complexes can explain a multitude of DNA folding patterns detected in chromosomes of diverse organisms including bacteria (Yatskevich *et al*., 2019). DNA loop extrusion has been observed with purified SMC complexes (cohesin and condensin) in single-molecule experiments (Davidson et al., 2019; Ganji et al., 2018; Kim et al., 2019). SMC motors move in large steps along DNA, in one or both directions, are able to bypass obstacles on the DNA relatively easily, but readily stall when the tension in the translocated DNA or to-be-translocated DNA increases. These unique features are explained for example by the ‘DNA segment capture’ model (also called ‘hold-and-feed’ model) where a small loop of DNA is captured between open SMC arms to be then fused with the previously captured DNA loop as the SMC arms close (Diebold-Durand et al., 2017; Nomidis et al., 2022). How DNA loop extrusion may promote the specific elimination of selfish DNA is not clear. Here we have first characterized the DNA substrate requirements of Wadjet systems in *B. subtilis* and *E. coli* indicating a limit in target DNA size and an exclusive preference for circular DNA. We then reconstituted an ATP-dependent enzymatic reaction for a selected purified JetABCD complex, leading to cleavage of several forms of circular plasmid DNA, but not of linear derivatives. The structure of its dimeric core complex was solved by cryo-electron microscopy (cryo-EM), elucidating a novel dimer-of-dimers configuration. Finally, we propose a hypothesis for the specific recognition of plasmid DNA by the self-collision of two DNA motor units of a JetABCD complex upon completion of extrusion of the extrachromosomal circular DNA molecule, eventually leading to DNA cleavage by the activation of an endonuclease function.

## Results

### JetABCD systems restrict circular plasmids in *B. subtilis* regardless of replication type and DNA uptake route

To characterize the substrate requirements of Wadjet and to uncover its limitations, we cloned three JetABCD operons from *Bacillus* species (*B. cereus, B. vireti* and *B. thuringiensis*) under the constitutive P*smc* promoter and integrated them at the *amyE* locus of the *B. subtilis* chromosome (Figure 1A) (Doron *et al*., 2018). The resulting strains were no longer able to be transformed with an 8.3 kb theta-replicating plasmid, pHCMC05, indicating hindrance of DNA uptake or plasmid maintenance by JetABCD as previously reported (Doron *et al*., 2018). Another plasmid, pC194, which undergoes rolling circle replication, did not yield colonies either when transformed into the JetABCD-harboring strains (Figure 1B). This was reverted when a conserved glutamic acid residue of the JetD TOPRIM domain was mutated to alanine (Figure 1B) (Doron *et al*., 2018). The lack of transformation is not due to defective natural competence as transformants still arose when the antibiotic resistant marker was integrated in the host chromosome rather than encoded on a plasmid (Figure S1A). Moreover, a larger 65 kb conjugative plasmid, pLS20, which is naturally resident in some *B. subtilis* strains, failed to transfer itself into these recipients (Figure 1C) (Itaya et al., 2006). The findings rule out the possibilities that the mode of plasmid replication or the route of DNA uptake are determinants for plasmid restriction by JetABCD.

### Plasmid restriction by JetABCD is sequence independent but limited by DNA size

We next wondered whether endogenously derived DNA would also be targeted by JetABCD. To test this, we excised DNA circles from the chromosome by homologous recombination. Briefly, we integrated two cassettes containing partially overlapping fragments of a kanamycin resistance gene (*neoR*) in direct orientation into the chromosome (Wu and Errington, 2002). Sporadic intramolecular homologous recombination events give rise to extrachromosomal circular DNA containing full-length *neoR*, a plasmid-derived replication origin as well as several essential genes, leading to loss of viability in case of its restriction by JetABCD (Figures 1D, S1B). Recombinants were selected by growth on agar plates supplemented with kanamycin. In cells expressing JetABCD, we did not obtain any kanamycin-resistant colonies when flanks were integrated at a distance of 10 kb, indicating that the excised 10 kb circular DNA was efficiently restricted (Figure 1E). Similar results were observed with a 35 kb spacing of the DNA flanks. However, when the flanks were placed at larger distances (from 57 kb to 85 kb), kanamycin-resistant colonies were observed with a higher rate (Figure 1E). This suggests that plasmid restriction by JetABCD is efficient on smaller plasmids, and that the size threshold above which restriction wanes appears to be in the range of fifty to hundred kilobases. As JetABCD can restrict chromosome-DNA-derived DNA circles, we conclude it does not discriminate DNA molecules based on DNA sequence or DNA modification.

### A linear plasmid evades restriction by *E. coli* JetABCD *in vivo*

SMC complexes are thought to extrude DNA loops to carry out their cellular function. We hypothesized that linear DNA would evade restriction as loop-extruding JetABCD complexes may dislodge from DNA at the ends. No linear plasmids have been described for *B. subtilis*. We thus established plasmid restriction in *E. coli*, which can host pJAZZ, a 12.3 kb linear plasmid with DNA hairpin telomeres derived from *E. coli* phage N15 (Figure 2A) (Ravin, 2011). We recovered a JetABCD operon from environmental *E. coli* strain GF4-3 (Foster-Nyarko et al., 2021) which has limited sequence homology and a shared operon organization to the counterpart from *B. cereus* (Figure 2B). The operon was placed under an arabinose-inducible promoter (P*bad*) and integrated into the *E. coli* K12 chromosome near the *glmS* locus by mini-Tn7 transposon-based insertion (Figueroa-Cuilan et al., 2016).

**Figure 2.**
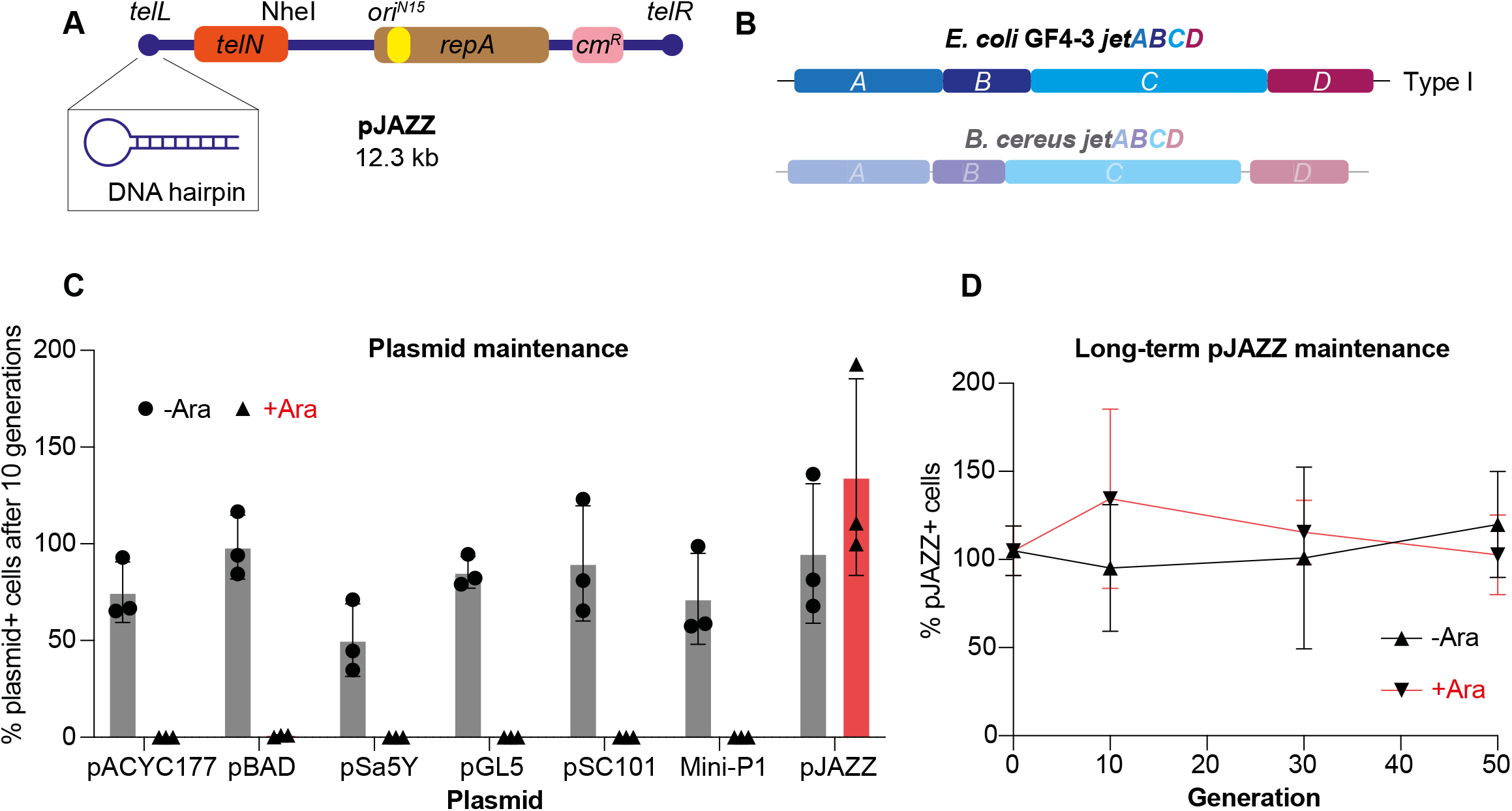
A linear plasmid can evade restriction by *E. coli* JetABCD. **A)** Schematic of pJAZZ with its DNA hairpin end depicted in the blowout. **B)** Operon organization for the GF4-3 *E. coli* JetABCD. For comparison the organization of the *Bc* system is also shown (Figure 1A). **C)** Graph showing the percentage of plasmid-containing *E. coli* cells (‘plasmid+’) in a cell population after ten generations without selection, with/without JetABCD induction by arabinose addition. Means and standard deviations from three independent experiments are shown. **D)** Graph showing the percentage of pJAZZ-containing *E. coli* cells in a population for up to 50 generations without selection, with/without JetABCD induction by arabinose addition. Means and standard deviations from three independent experiments are shown. See also Figure S2.

Plasmid restriction appeared less robust in *E. coli* as transformations yielded normal numbers of colonies despite induction with arabinose before or after transformation. However, the stability of plasmids in cells grown for multiple generation in liquid culture in the absence of antibiotic selection was markedly reduced when *jetABCD* expression was induced with arabinose. Six tested circular plasmids with different replication origins (Table S3) were essentially eliminated from the cultures after ten generations of growth without selection (Figure 2C). As expected, plasmid loss was neither observed in *E. coli* expressing the ATPase-dead mutant JetC(E1022Q, ‘EQ’), nor JetD(E248A) (equivalent to E226A, E295A and E298A mutations of *Bacillus cereus, vireti*, and *thuringiensis* JetD, respectively of Figure 1B) (Figure S2A). pJAZZ, however, remained remarkably stable even after 50 generations of growth (Figures 2C, 2D), suggesting that linear DNA can indeed efficiently evade plasmid targeting by JetABCD. Of note, no obvious effects on plaque formation upon infection with several tested *E. coli* phages (T4, T6, T7, P1, and lambda) was observed (Figure S2B) as previously found for *B. subtilis* (Doron *et al*., 2018), implying that lytic cycles are not affected by JetABCD. We conclude that JetABCD targets smaller-sized circular DNA molecules but not linear or larger DNA molecules, including the host chromosome(s).

### Purified JetABCD cleaves circular but not linear DNA molecules

To better understand how JetABCD recognizes a smaller circular DNA molecule and to study its fate, we next aimed to reconstitute the activity of a JetABCD holo-complex by recombinant expression of subunits in *E. coli*. When compared to the three *Bacillus* variants, JetABCD proteins from *E. coli* strain GF4-3 supported better protein expression and purification. It was thus chosen for detailed analysis. Plasmids harboring wild-type (but not mutant) versions of all JetABCD components were poorly co-transformed into *E. coli* strains, presumably due to JetABCD restricting its own expression plasmids. We thus resorted to expressing and purifying JetD separately from the other components. Affinity chromatography, ion exchange chromatography, and/or size exclusion chromatography (SEC) were used to produce preparations of untagged JetD and of JetABC complexes carrying a 10His-Twin-Strep-tag on the N-terminus of JetA. To reconstitute the holo-complex, JetABC was mixed with JetD and analyzed by SEC showing that the four subunits co-eluted from the column in approximately stoichiometric amounts (Figure 3A, S3A). When performing initial DNA binding assays by electrophoretic mobility shift experiments, we noticed that the gel mobility of a large test plasmid was surprisingly increased upon pre-incubation with JetABCD in our agarose gel running system. The increased mobility approximately corresponded to the mobility of plasmid DNA linearized by a restriction endonuclease, indicating that the plasmid was cleaved by JetABCD. Normal mobility was not recovered by protein denaturation using SDS addition and heat treatment, supporting the idea of a covalent modification of DNA. Protein denaturation was used routinely prior to gel analysis in the following experiments.

**Figure 3.**
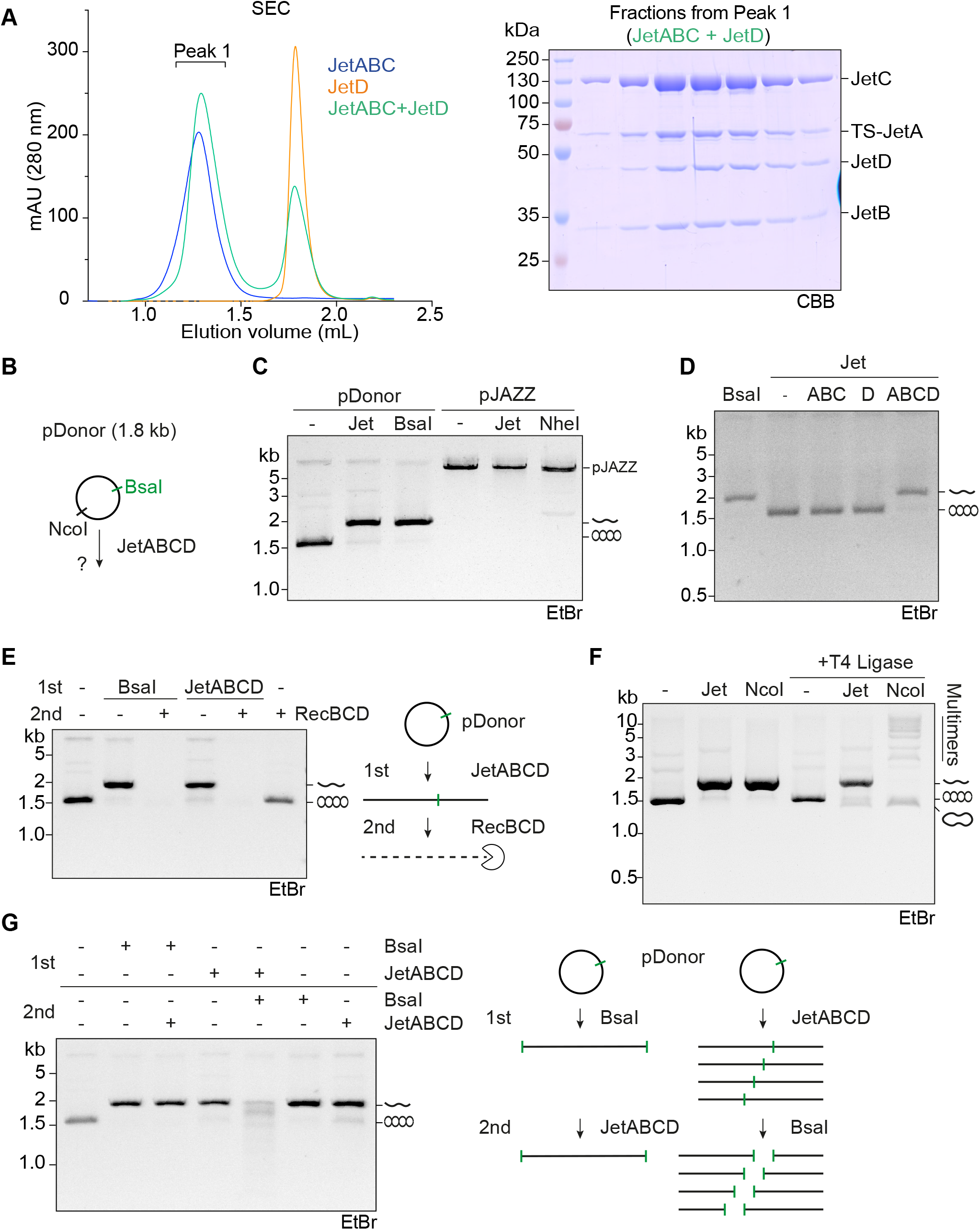
Purified JetABCD cleaves circular but not linear DNA molecules. **A)** Reconstitution of JetABCD. Left panel: Elution profiles for analytical SEC of JetABC and JetD. Right: Peak fractions of the JetABCD elution analysed by SDS-PAGE and Coomassie Brilliant Blue (‘CBB’) staining. ‘TS’, 10His-Twin-Strep-Tag. **B)** Schematic of circular test plasmid pDonor used for cleavage experiments. See Figure S3C for more detailed schematic. **C)** Plasmid DNA cleavage reactions with JetABCD. JetABCD was reconstituted by mixing 12.5 nM of JetABC dimer-of-dimers and 25 nM of JetD (final concentrations). pDonor (8.5 nM) or pJAZZ (3.75 nM) were mixed with ATP (1 mM) and incubated with JetABCD for 15 minutes at 37°C. The reaction was stopped by SDS/heat treatment and the products analyzed by ethidium bromide (‘EtBr’)- agarose gel electrophoresis. **D)** Plasmid DNA cleavage assay with JetABC, JetD, and JetABCD. **E)** Treatment of the JetABCD cleavage products with the exonuclease RecBCD. Note that Streptavidin-blockage of DNA ends on a linear DNA molecule hindered degradation by RecBCD (Figure S3D). **F)** Ligation of JetABCD cleavage products with T4 ligase. Only a small fraction of the cleaved product is converted to covalently closed circular DNA (relaxed DNA). **G)** Post-treatment (but not pre-treatment) of JetABCD cleavage products with the restriction endonuclease BsaI results in a DNA smear. All gels shown are representative examples from at least two independent experiments. See also Figure S3.

Cleavage activity by JetABCD was observed with a variety of circular plasmid DNA substrates but not with linear pJAZZ (Figures 3B, 3C, S3B), mirroring what we found for plasmid restriction *in vivo* (Figure 2C). We proceeded with a 1.9 kb test circular plasmid, denoted as pDonor, for subsequent experimentation (Taschner et al., 2021) (Figure S3C). JetABC and JetD alone did not show activity, indicating that all four subunits are needed for cleavage (Figure 3D).

To characterize the nature of the DNA ends generated by JetABCD cleavage, we first incubated the JetABCD cleavage product with the exonuclease RecBCD (Simon et al., 2016). This resulted in the complete elimination of the JetABCD-cleaved DNA species but not of uncleaved plasmid DNA (Figures 3E and S3D). We also treated JetABCD-cleaved DNA with proteinase K prior to gel analysis to remove any covalently attached polypeptides. No obvious alteration in the gel running behavior was detected (Figure S3E). These findings strongly suggest that JetABCD-generated DNA ends are not blocked by proteins and indicate that JetABCD may hydrolyze both strands of the DNA double helix to generate free DNA ends. Accordingly, JetABCD is not a suicide enzyme that remains covalently associated with cleaved DNA (like the distantly related TOPRIM containing protein Spo11). Notably, T4 ligase treatment only poorly but detectably re-circularized the JetABCD cleavage product, implying that DNA cleavage generated compatible ends with 5’ phosphates on some plasmid molecules (Figure 3F). No plasmid multimers were detected after ligation (as predominantly seen after NcoI digest) hinting at a preference for intra-molecular ligation. Possibly, DNA ends generated on different DNA molecules are incompatible, for example due to differences in the base sequence of the overhangs.

SMC complexes (on their own) generally do not show sequence specificity in DNA binding. To test whether DNA cleavage occurs with preference for certain sequence(s) or not, we combined JetABCD incubation with pre- or post-treatment by the BsaI restriction endonuclease which cuts at a defined position. As expected for random DNA cleavage by JetABCD, subsequent treatment with the restriction endonuclease converted the linear band species into a smear of shorter DNA fragments (Figure 3G). The smear was not completely uniform, indicating that JetABCD may have slight target site preferences. Importantly, treatment of plasmid DNA with the restriction endonuclease prior to incubation with JetABCD produced full-length linear DNA rather than a smear (Figure 3G), as expected from the specificity of JetABCD for circular DNA.

### Strict requirements for DNA cleavage by JetABCD

We next investigated the requirements for cleavage activity by JetABCD. No DNA cleavage was observed without ATP (Figure 4A) indicating that DNA cleavage depends on the SMC ATPase. To test this more directly, we produced an ATPase mutant complex harboring JetC(EQ). Equivalent Walker B motif mutations are widely used in ABC-type ATPases to prevent ATP hydrolysis but permit normal ATP binding and ATP-head engagement. As expected, JetC(EQ) was unable to support ATP hydrolysis despite being well expressed and efficiently producing JetABCD holo-complexes (Figure S4A). It also failed to promote ATP-dependent DNA cleavage (Figure 4A). This implies that ATP binding, head engagement, and ATP hydrolysis are all needed (at least once) for DNA cleavage, being consistent with the idea that DNA cleavage requires a DNA-motor function of JetABCD. Apart from the Mg^2+^ that we normally supply together with ATP in the reaction buffer (HEPES pH 7.5 10 mM, KOAc 150 mM, MgCl_2_ 5 mM, ATP 1 mM, TCEP 1 mM), no addition of further divalent cations was needed for DNA cleavage, implying that the putative endonuclease works with Mg^2+^ or without divalent cation.

**Figure 4.**
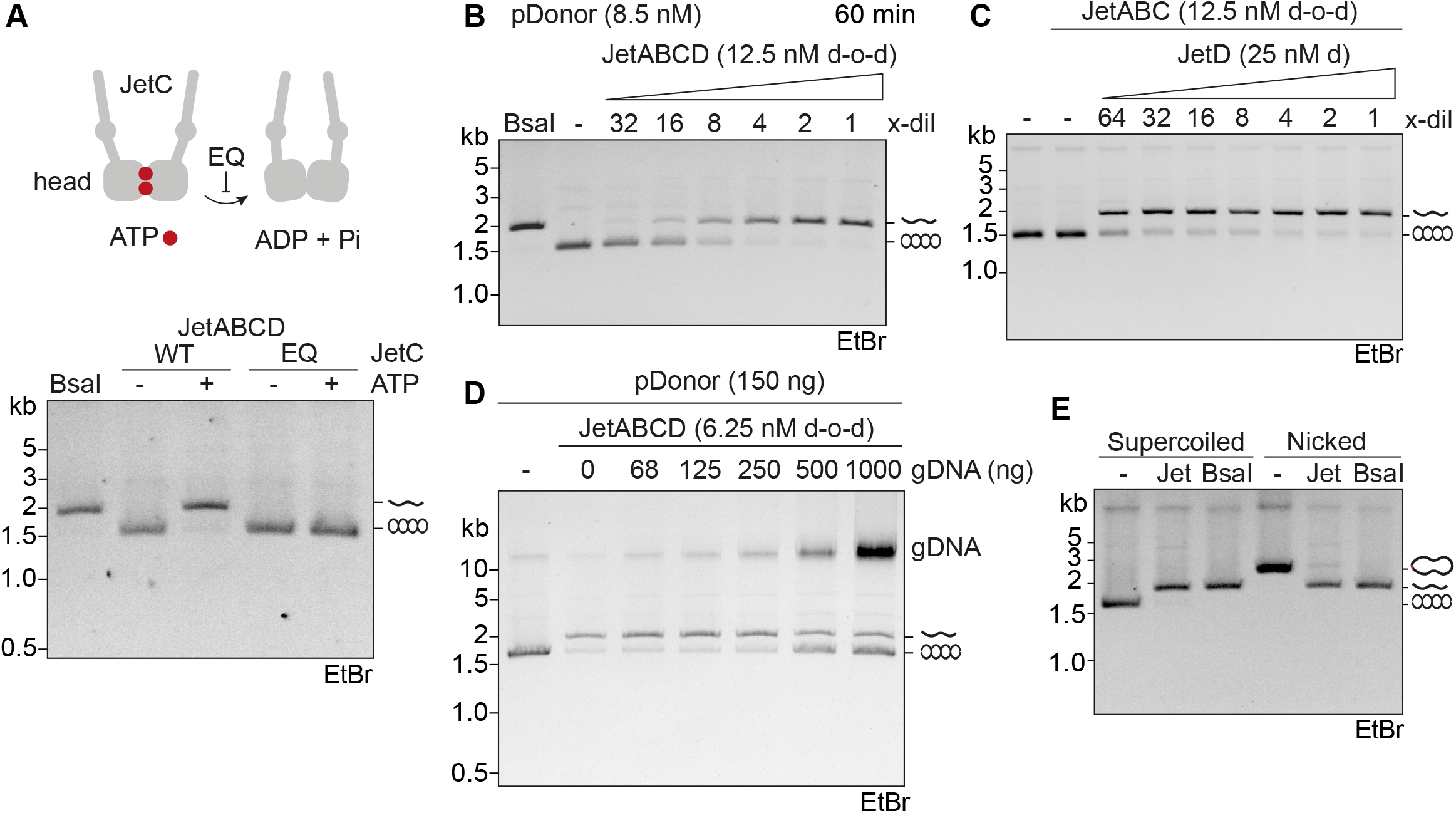
Requirements for plasmid DNA cleavage by JetABCD. **A)** Plasmid DNA cleavage by JetABCD in presence or absence of ATP with JetC(WT) and JetC(EQ) mutant complexes. **B)** Plasmid DNA cleavage with limiting amounts of JetABCD; incubation for 60 min at 37°C. **C)** Plasmid DNA cleavage with limiting amounts of JetD. ‘d’, dimer. **D)** Competition assay for plasmid cleavage by the addition of increasing amounts (up to 7-fold excess over plasmid DNA) of linear genomic DNA (gDNA) prepared from *B. subtilis*. Note that the large gDNA fragment migrate near the top of the gel. **E)** JetABCD cleavage assays with supercoiled and nicked forms of pDonor plasmid. Plasmids were pre-treated with the nicking endonuclease Nt.BspQI (‘Nicked’) or mock-treated (‘Supercoiled’). After heat inactivation and tenfold dilution into assay buffer, the samples were incubated with JetABCD. JetABC(D) concentrations are denoted for the dimer-of-dimers (d-o-d) complex. See also Figure S4.

Titration experiments with decreasing amounts of JetABCD indicated that the turnover number is relatively low. After prolonged periods of incubation (60 min), a given JetABCD complex cleaves more than one plasmid (Figures 4B, S4B). In contrast, somewhat lower, sub-stoichiometric amounts of JetD were able to support efficient cleavage, implying that JetD is not involved in the rate-limiting step of the process. JetD may not be bound to JetABC at all times and is able to be recruited by JetABC for the processing of a substrate (Figure 4C). Competition experiments using excess of linear DNA (genomic DNA, ‘gDNA’) however reduced the DNA cleavage activity, implying that JetABC also binds to linear DNA but without processing it (Figure 4D).

We did not observe DNA cleavage when using JetD(E248A) mutant instead of wild-type JetD (Figure S4E) (Doron et al, 2018). Moreover, DNA cleavage by JetABCD(WT) was inhibited by an excess of JetD(E248A) (Figure S4E), suggesting that JetD(E248A) can outcompete JetD binding to JetABC.

We next investigated how plasmid DNA topology impacts cleavage by JetABCD. In proliferating bacterial cells, DNA is typically underwound (‘negatively supercoiled’; having a deficit in helical turns) due to the combined action of transcription and topoisomerases (Dorman, 2019). A relaxed conformation of plasmid DNA can be obtained *in vitro* by breaking of one of the two DNA strands (‘nicked’ DNA) by a nicking endonuclease. Re-ligation of the DNA nick results in a covalently closed circular form that is now devoid of underwinding (‘relaxed’). Four forms of plasmid DNA (supercoiled, linear, relaxed, nicked) can be separated by agarose gel electrophoresis in the presence of ethidium bromide (EtBr), whose DNA intercalation leads to local DNA underwinding and, in covalently closed circular molecules, to compensatory positive supercoiling. We found that neither the nicked (by Nt.BspQI) nor the relaxed form of pDonor evaded DNA cleavage by JetABCD (Figures 4E, S4C). Like cleavage of supercoiled pDonor DNA, cleavage of the nicked form required ATP, JetD as well as JetABC (Figure S4D). These results suggest that the helical topology of DNA is not a critical determinant of cleavage. Moreover, the building up of helical stress by an SMC-motor action is probably not required for plasmid restriction, at least *in vitro*, as helical stress is expected to be dissipated rapidly on nicked DNA (unless being constrained by JetABCD). Of note, we find that the presence of the nick does not obviously alter target site preference of JetABCD (comparing Figure 3G and S4D).

Altogether, we conclude that plasmid restriction is likely mediated by DNA cleavage. JetABCD cleaves DNA with a high specificity for circular plasmid DNA of multiple forms, which is likely governed by a SMC ATPase DNA-size-and-shape-sensing mechanism and the ensuing activation of a JetD-endonuclease reaction.

### Cryo-EM structure of the JetABC core reveals a novel DNA-motor dimer arrangement

To better understand how the SMC motor may instruct the DNA cleavage reaction, we investigated the structure of (DNA-free) JetABCD by cryo-EM. The reconstituted *E. coli* JetABCD complex (mixed with ATP) was well suited for cryo-grid preparation, yielding grids with well-dispersed particles. We collected about 6000 micrographs on a Titan Krios, automatically picked more than 600,000 particles and obtained 2D class averages (Figure S5A, S5B).

In the resulting cryo-EM map, we observed a compact dimer-of-dimers configuration with two JetC motor units intimately held together by a central JetAB subcomplex. In the structure, the JetC dimers are oriented relative to each other in a narrow angle (roughly 30°) and with opposing orientations (Figure 5A). This V-shaped ‘closed’ configuration contrasts a much more ‘open’ I-shaped configuration observed for MukBEF where two motors are facing in the same direction (Burmann *et al*., 2021) (Figure 5D). The alternative dimer-of-dimers configuration is governed by distinct contacts formed between N-terminal JetA sequences and the JetC proteins (discussed further below). In isolation, all visible domains resemble corresponding parts from other SMC complexes, particularly MukBEF (Figure S6A). No obvious density was observed for the more distal JetC coiled coils, presumably due to intrinsic structural flexibility, and for JetD. Notably, the hinge is also not visible in the map, suggesting that the hinge does not come into the vicinity of the core complex as seen in several other SMC complexes such as MukBEF, cohesin, and condensin but not in Smc-ScpAB and Smc5/6 (Burmann et al., 2019; Lee et al., 2020; Soh et al., 2015; Taschner *et al*., 2021).

**Figure 5.**
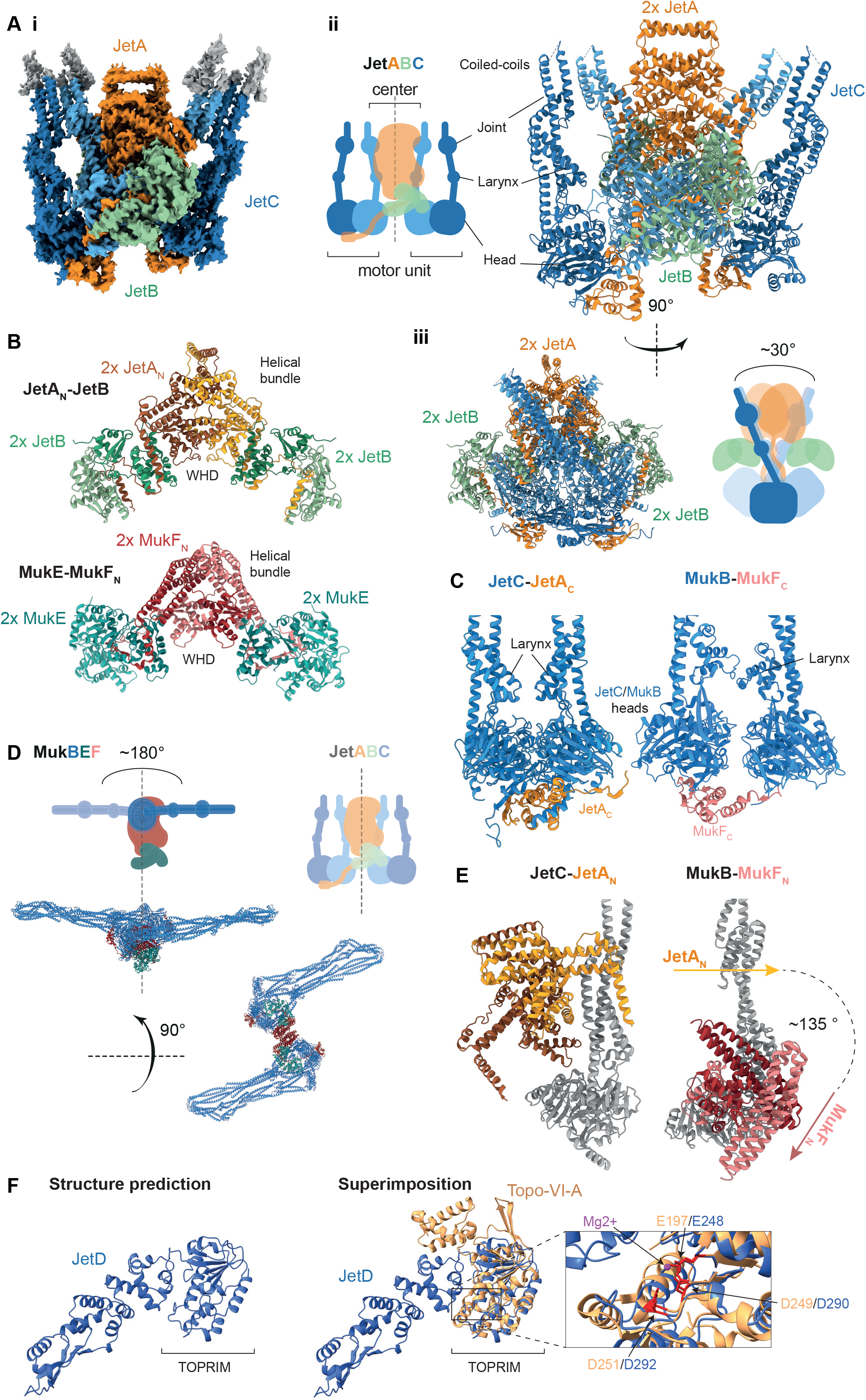
Cryo-EM structure of the JetABC core reveals a novel DNA-motor dimer arrangement. **A) i**, Density map of JetABC (global resolution: 3Å) showing its dimer-of-dimers architecture with JetA subunits in yellow and orange colours, JetB subunits in green colours and JetC in blue colours. **ii**, Pseudo-atomic model of the core of the JetABC dimer-of-dimers with schematic in corresponding colours. **iii**, a side view (90-degree-rotated along the y-axis) showing a V-shaped motor unit arrangement. **B)** Comparison of JetA_N_ dimer (top panel) and MukF_N_ dimer (bottom panel, PDB: 7nz4). Side views of JetA and MukF bound to two dimers of JetB and MukE, respectively, are shown. See Figure S6B for top view. **C)** Comparison of JetC-JetA_C_ interaction with MukB-MukF_C_ (PDB: 7nz4). **D)** Top: Schematic and structures of MukBEF oriented by aligning MukEF like JetAB in Figure 5A. **E)** Comparison of the JetA_N_ to JetC and MukF_N_ to MukB association revealing a large angular change and shift in JetC position and JetA binding subunit (front in JetA; back in MukF). **F)** Alphafold structure predictions of JetD and comparison with topoisomerase VI-A subunit. From left to right: **i**, JetD predicted by AlphaFold, **ii**, Superimposition with TopoVIA crystal structure (PDB: 1d3y) guided by the shared TOPRIM domain. Inset: Note that residues coordinating Mg2+ ions in the mjTopoVIA TOPRIM domain (E197, D249, and D251) are conserved in JetD (E248, D290, and D292). See also Figure S5 for the cryo-EM pipeline and S6 for more structural comparisons between JetABC and MukBEF.

The JetABC dimer-of-dimers is comprised of three substructures: a JetAB center and two peripheral JetC motor units (Figure 5A). These substructures individually resemble the matching substructures of MukBEF (Figure 5B, C). The central JetAB subcomplex is formed from a dimer of JetA N-terminal sequences (JetA_N_) as well as four JetB kite proteins. Like in MukF_N_, JetA_N_ folds into a winged-helix domain (WHD) and a helical bundle, which together form a tight homodimer (Burmann *et al*., 2021; Fennell-Fezzie *et al*., 2005). The JetB proteins form a canonical homodimer and bind to the otherwise unstructured JetA middle segment (Palecek and Gruber, 2015). The motor units are made up of a JetC SMC dimer bound via one of the two head domains, the κ-JetC head, to the C-terminal JetA WHD (Figure 5C). The heads appear in a juxtaposed, ATP-non-engaged conformation (despite the presence of ATP during grid preparation) (Burmann *et al*., 2021; Diebold-Durand *et al*., 2017). The overall architecture of the JetABCD structure deviates from MukBEF (Figure 5D) by a different association of the central unit and the two motor units. In the cryo-EM structure, the JetA helical bundle associates with the κ-JetC head-proximal coiled coil helices in a perpendicular manner (Figure 5E) with additional contacts being provided by the JetC coiled coil and the JetA WHDs. In MukBEF, however, the MukF helical bundle is bound to the MukB head at a lower position and oriented almost in parallel to the MukB coiled coil (Figure 5E). This large angular change results in an almost 180° re-configuration of the two motor units in the dimer-of-dimers (Figure 5D). In the closed configuration the JetB kite subunits are nevertheless positioned close to the head domains, possibly indicating that they can support DNA clamping after minor reconfiguration. Thus, the closed configuration may conceivably be competent for DNA translocation and loop extrusion. Curiously, a given JetA subunit connects a protomer from one JetC dimer to a protomer from the other JetC dimer, rather than forming the tripartite rings seen in other SMC complexes (Figure 5A). If true, this has potential implications for DNA loading as the respective ring interface has been implicated in ring opening for DNA loading in some SMC complexes. DNA loop extrusion however might not be affected by the alternative subunit connectivity, assuming it does not require gate opening.

We also probed JetA-JetC associations by structural predictions with AlphaFold. Using relevant fragments of JetA and JetC as input sequences, the predictions readily returned structural models that more closely resemble the MukBEF architecture than the JetABC cryo-EM structure (Figure S6C). This possibly suggests that the JetABC complex is also able to adopt a MukBEF-like open configuration and if so, it is conceivable that the open configuration supports DNA loop extrusion (as expected for MukBEF) while the closed configuration confers DNA cleavage. Alternatively, the AlphaFold prediction might be an artefact due to a bias towards the open conformation caused by the presence MukBEF entries in the learning dataset. The significance of these forms of JetABCD thus remains to be established.

Since JetD was absent from the cryo-EM structure (despite being present during cryo-grid preparation) we modelled it by AlphaFold. The JetD AlphaFold model displayed highest similarity with the TOPRIM domain of the DNA cleavage subunit A of DNA topoisomerase VI (Figure 5F) (Corbett et al., 2007; Nichols et al., 1999). On the JetD TOPRIM domain, we readily recognized the DxD motif residues (D290 and D292) and a conserved glutamate (E248) that in topoisomerase VI coordinate a Mg^2+^ ion for DNA binding and cleavage of the scissile bond (Nichols *et al*., 1999). The fold of the adjacent domain (a WHD in topo VIA) showed more limited similarities; we failed to identify a residue that is equivalent to the catalytic tyrosine undergoing transesterification in topoisomerase VI. The apparent absence of the catalytic tyrosine is consistent with the notion that JetABCD hydrolyses the DNA phosphodiester backbone (rather than undergoing transesterification). Other conserved JetD residues (such as K172, R182, E362, and E364) (PFAM: DUF2220, DUF3322) likely contribute to catalysis instead.

## Discussion

Self-nonself discrimination is a ubiquitous challenge in biology. We show here that JetABCD specifically recognizes plasmids based on their circular nature and limited size. Canonical SMC complexes support a multitude of chromosome-related processes by DNA loop extrusion. The related Wadjet system putatively harnesses DNA loop extrusion to specifically recognize plasmids for restriction. Our finding that DNA cleavage requires JetC ATP hydrolysis corroborates the idea that plasmid restriction rely on a SMC DNA motor function. Plasmid restriction by JetABCD eliminates DNA actively by DNA cleavage rather than by DNA passivation through sequestration or silencing. However, DNA cleavage is a highly dangerous activity that requires strict control. Here, we propose a mechanism for extrachromosomal DNA recognition and elimination by JetABCD with possible implications for viral restriction by Smc5/6.

### A model for specific elimination of smaller circular DNA molecules

We show here that JetABCD discriminates DNA molecules predominately on the basis of their circular nature and limited size, apparently regardless of DNA helical topology. To explain how JetABCD efficiently targets circular plasmids without endangering genome integrity by generating DNA breaks on the chromosome, we propose that DNA cleavage only occurs when a single JetABC(D) complex completes the extrusion of an entire plasmid molecule. In this situation, the two motor units of the respective JetABCD complex are expected to approach one another (Figure 6A, panel i), eventually resulting in the stalling of DNA translocation. The stalling in turn may lead to the buildup of tension within the motor complex and/or the remainder of the DNA possibly generating the active DNA cleavage unit (Figure 6A). The tension may thus serve as signal for the activation of the DNA cleavage reaction, for example by allowing the engagement of two JetD subunits that are otherwise kept apart by being bound to opposite motor units or by the reconfiguration of the dimer-of-dimers architecture. The tension however might not critically rely on DNA over- or underwinding since we find that nicked plasmid DNA is targeted by JetABCD almost as efficiently as negatively supercoiled DNA (Figure 4E, S4D).

**Figure 6.**
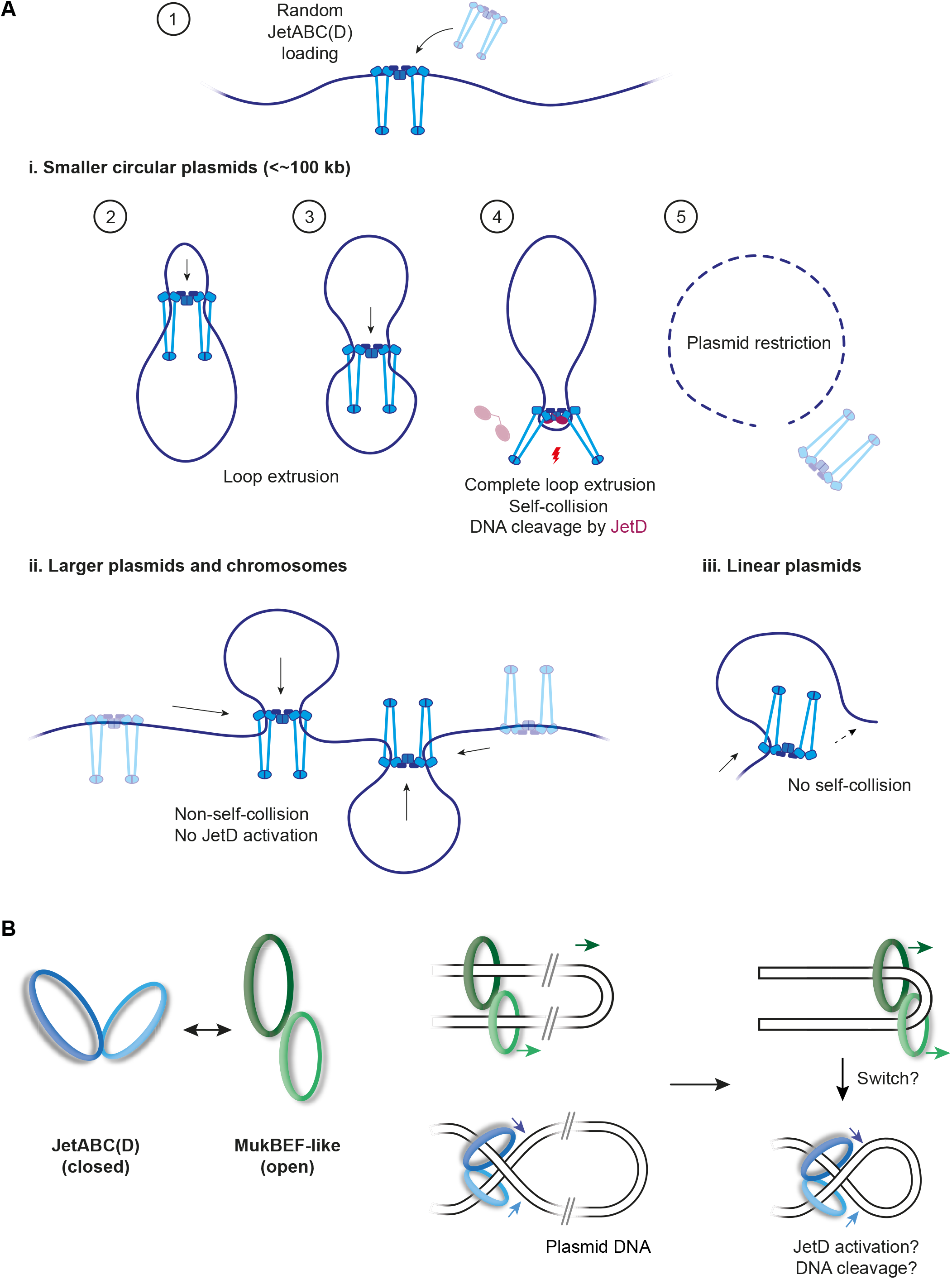
Model of plasmid restriction by JetABCD. **A)** JetABCD randomly binds to its target DNA (top panel), with the following consequences depending on DNA type: **i)** Smaller plasmids: JetABC self-collision occurs when all DNA has been extruded, resulting in DNA tension that could trigger conformational changes, resulting in JetD recruitment or activation and thereby DNA cleavage. **ii)** Larger plasmids and chromosomes: Complete loop extrusion is less/not permissive, no JetABC self-collision occurs and therefore restriction by cleavage cannot occur. **iii)** Linear plasmids: complete loop extrusion and JetABC self-collision cannot occur as the complex would be dislodged from DNA.**B)** Schematic of how the contrasting MukBEF (green) and JetABCD (blue) motor unit arrangements results in different conformations of extruding plasmid DNA loops. For JetABCD, two DNA double-helices would thread through in an overlapping, cross-shaped manner while they would be parallel to each other for MukBEF. Just prior to loop extrusion completion, JetABCD would create a teardrop-shaped DNA loop, the shape or tension of which could serve as a signal for downstream JetD-mediated cleavage.

On the chromosome, JetABCD complexes are likely to meet other complexes or to stochastically dissociate from DNA prior to completing extrusion due to the larger size of chromosomal DNA (Figure 6A, panel ii). Linear DNA molecules also hinder the generation of self-stalling motor complexes due to the dislodging of motor units at the DNA ends (Figure 6A, panel iii). A prediction from this hypothesis is that elevated levels of JetABCD may lower the target size limit due to an increased frequency of collisions between JetABCD complexes; preliminary results however did not support this notion (Figure S1C), possibly indicating that target size limitation might be mostly governed by a finite processivity or chromosome residence time.

This simple hypothesis—while being entirely speculative—readily explains self-nonself discrimination. It may equally well apply to viral recognition by Smc5/6 even if DNA cleavage might be replaced by a more passive mechanism for DNA inactivation such as the silencing of gene expression from the circular viral genome or a block in DNA replication.

### Differences with known SMC complexes

Our structure of the JetABCD core provides first hints at the underlying structural mechanisms. As described above, the two motor units are arranged distinctly from MukBEF. The V-shaped arrangement of the motor units will lead to unique organization of DNA with potential implication for one or more steps of the activity cycle, *i*.*e*. for DNA loading, translocation, and restriction.

The alternative geometry has potential implications for the presumed DNA motor activity of these SMC complexes. The open configuration of the two motors in MukBEF allows them to reel DNA in from the same side, rather than from opposite sides as would be expected from a stacked motor geometry in JetABCD (Figure 6B). We speculate that the closed configuration may be particularly relevant during the final stages of DNA restriction, when the two motor units approach each other, then stall, leading to tension in the remaining DNA. A reconfiguration from an initially open to the closed configuration upon the buildup of DNA tension is also possible; it might serve to activate DNA cleavage. Assuming that the DNA segment capture model (Diebold-Durand *et al*., 2017; Nomidis *et al*., 2022) applies to JetABCD, we can predict the directionality of DNA transport by the JetABCD dimer (Figure 6B). Intriguingly, during termination, the closed configuration would lead to the formation of a small circle- or teardrop-shaped DNA in the remaining DNA, which conceivably may act as signal for DNA cleavage activation, for example, by generating a higher-affinity binding site for a JetD dimer (Figure 6B, right panels).

Currently, we cannot exclude the possibility that the closed configuration is a resting state of JetABCD or even an artefact of recombinant protein production. It will be exciting to determine whether the V-shaped organization of SMC motors is a unique feature of complexes with restriction functions or also found in DNA loop extrusion complexes. Solving the structures of other JetABCD systems, particularly of those with diverged sequences, will be instrumental. The comparison with other SMC complexes is important to conclusively determine whether the altered motor geometry is crucial for plasmid restriction or a byproduct of divergent evolution. At the same time, it is important to detect these configurations in solution at different steps of the restriction process.

It is noteworthy that another endonuclease also displays an SMC-like architecture. Unlike JetABCD however, Rad50/Mre11 specifically recognizes linear DNA molecules and cleaves at the DNA ends to initiate DNA repair (Kashammer et al., 2019). Remarkably, machines sensing the presence and absence of DNA ends, respectively, share a common ancestry.

### Evolutionary implications

Defense systems generally exhibit higher levels of sequence divergence, presumably due to elevated selection pressures exerted by rapidly evolving selfish DNA elements. The relatively high levels of divergence observed in JetABCD presumably reflects its history as defense system. It will be exciting to identify plasmids that evade restriction by JetABCD, to uncover the underlying mechanisms, and to reveal how JetABCD may adapt to counteract the evasion mechanism. MukBEF shows similar sequence divergence from the more canonical SMC complexes, despite its involvement in a housekeeping function, possibly indicating that it also underlies high selection pressure or more likely that it emerged from an ancestral JetABCD-like immunity system. Our structure and sequence analysis reveals features of JetABCD that it shares exclusively with MukBEF (the dimer-of-dimers configuration, reduced hinge domain size, elbow folding albeit at a different angle), others that are more closely related to Smc-ScpAB (arms with fewer coiled coil deviations, simpler joint, and some that are intermediary (such as a simplified version of the MukBEF larynx). These observations further support the idea that MukBEF has emerged from a JetABCD-like ancestor. If so, then the mechanisms of DNA extrusion by the respective defense and chromosome organization systems must be principally equivalent and interchangeable. Conceivably, JetABCD systems may even contribute to chromosome organization or segregation while scanning cellular DNA for the presence of foreign elements. They may thus represent dual function machines promoting chromosome maintenance and plasmid DNA anti-maintenance all at the same time.

### Possible applications

Like other DNA-based immunity systems including R/M and CRISPR/Cas, the unique DNA cleavage activity of JetABCD may serve as powerful tool in research and biotechnology. We envision its implication in the elimination of plasmids from DNA preparations, the identification or isolation of DNA circles for diagnostic purposes (e.g. in cancer patients), the mutagenesis of DNA (*e*.*g*. by DNA insertion) at randomized positions for screening purposes. Finally, its application for the specific elimination of extrachromosomal circular DNA (‘plasmid curing’) from cells or from microbes are in principle conceivable for treatments against viral infection, cancer, antibiotic resistance, and cellular aging (Kumar et al., 2020; Vrancianu et al., 2020).

## Acknowledgements

We are grateful to the Dubochet Centre for Imaging in Lausanne for cryo-grid preparation and cryo-imaging, to Kyle Muir for continuous help during initial cryo-EM data analysis, and to Michael Taschner for advice in recombinant protein production and purification. We thank David Adams and Melanie Blokesch for help and materials for *E. coli* genetic engineering and to Mark Pallen for kindly providing the *E. coli* strain GF4-3. This work was supported by the European Research Council (724482 to S.G.). HWL is supported by an EMBO Postdoctoral fellowship (ALTF 490-2021).

## Author contributions

Conceptualization, S.G., H.W.L., and F.R.H; Methodology, B.B., A.M., Y.L., H.W.L., and F.R.H; Investigation, H.W.L., and F.R.H; Writing – Original Draft, H.W.L., and S.G.; Writing – Review & Editing, all authors.; Funding Acquisition, S.G.; Resources, S.G.; Supervision, S.G.

## Declaration of Interests

The authors declare that they have no competing interests.

## Notes

### Competing Interest Statement

The authors have declared no competing interest.

