## Supplementary material for "DNA-measuring Wadjet SMC ATPases restrict smaller circular plasmids by DNA cleavage": Methods & Material and Supplementary Figures

***Star Methods***

#### Resource Availability

**Lead Contact**

**Materials Availability**

All DNA constructs and strains generated in this study are available from the lead contact without restriction.

**Data and Code Availability**

The cryo-EM data has been deposited at the PDB (8AS8) and EMBD (EMD-15609) data banks.

#### Experimental Model and Subject Details

**Strain construction in *B. subtilis***

The *B. subtilis* strains used in this work originate from either the 168 ED or 1A700 isolate. Natural competence was used to engineer strains at target loci by allelic replacement, as described in (Diebold-Durand et al., 2019). Strains were selected on nutrient agar plates under appropriate antibiotic selection. Genotypes were verified for single colony isolates by PCR and Sanger sequencing (Microsynth) as required. *jetABCD* operons from *B. cereus, B. vireti, B. thuringiensis* were synthesized from TWIST biosciences. A list of strains and genotypes is provided in Supplementary Table 1. The assignment of strains to each figure panel is listed in Supplementary Table 2.

**Transformation assay in *B. subtilis***

Transformation assays were performed with naturally competent *B. subtilis* cells. For pHCMC05, 250ng of plasmid DNA was used per assay. To introduce pC194, we used 100ng isolated genomic DNA from BSG4800. To introduce genomic DNA, we used 40ng of DNA from a strain carrying the appropriate chromosomal antibiotic resistance marker. In all assays, plates containing transformant colonies were imaged and colonies counted with ImageJ. All plasmids tested in this study are listed in Supplementary Table 3.

**Conjugation assay in *B. subtilis***

Donor and recipient strains were grown overnight in LB media with appropriate antibiotic selection. After back-dilution into fresh LB, cultures were allowed to grow until exponential-phase. 1 $OD_{600}$ unit of the donor and recipient cultures were harvested, mixed and filtered onto a 0.45 µm cellulose nitrate filter (Sartorius). Filters were transferred onto nutrient agar plates without selection and were incubated at 37°C for 5 hours. Cells were dislodged from the filter in LB, washed once in LB and resuspended in a final volume of 10 mL LB. 100 µL of the resuspension was plated on nutrient agar plates with antibiotic election for transconjugants (i.e. containing both donor and recipient resistance markers) and incubated overnight at 37°C. In all assays, plates containing transconjugant colonies were imaged and colonies counted with ImageJ.

**Chromosome excision assay**

Homologous recombination flank-containing strains were grown overnight in LB media supplemented with glucose (0.5 % (w/v) final) and the appropriate antibiotic at 30°C. They were back-diluted to fresh LB + 0.5% (w/v) glucose and allowed to grow until late exponential phase ($OD_{600}$=0.8) at 30°C. 200 µL of culture was harvested and plated on nutrient agar supplemented with kanamycin (2 µg/mL final). After overnight incubation at 37°C, plates containing kanamycin-resistant recombinants (i.e. harboring DNA circles) were imaged and colonies counted with ImageJ.

Where P*xyl-jetABCD* strains were used, cultures were grown in LB supplemented with xylose (0.5 % (w/v) final) for *jetABCD*-induced conditions and glucose (0.5% (w/v) final) otherwise. For plating, xylose-supplemented nutrient agar plates were used for the induced condition.

**Strain construction in *E. coli***

A mini-*Tn7* transposon carrying *araC* and GF4-3 *jetABCD* under the P*bad* promoter was integrated into a neutral *E. coli* chromosomal locus downstream of *glmS* by triparental mating, as previously described (Bao et al., 1991). Plasmids were introduced into electrocompetent *E. coli* cells by electroporation at 2.0 kV.

**Plasmid stability assay in *E. coli***

*E. coli* cultures were grown overnight at 37°C in LB media with appropriate antibiotic for plasmid selection. Each strain was back-diluted into LB at an $OD_{600}$ of 0.0025 either in the absence or presence of arabinose (0.02 % (w/v) final) and allowed to grow for approximately 10 generations. 200 µL of culture was harvested and serially diluted sevenfold (from $10^{-1}$ to $10^{-7}$) in PBS. For total cell number count, 5 µL of the dilutions from $10^{-4}$ to $10^{-7}$ were spotted onto nutrient agar plates in duplicate. For plasmid-containing cells, the same amount of all dilutions was spotted onto nutrient agar plates supplemented with the plasmid-selecting antibiotic. After overnight incubation at 37°C, colonies were counted and extrapolated based on the dilution degree to give an estimate of the total cell number and survivor count. To quantify plasmid+ cells, we calculated the percentage of plasmid+ cells over the total cell number. Datapoints greater than 100 % were sometimes obtained if the number of plasmid+ colony count was greater than total cell colony count.

**Phage protection assays**

Overnight cultures of *E. coli* strains were back-diluted LB supplemented with either 0.02 or 0.2 % (w/v) % arabinose and grown at 37 °C with shaking for 2 h. Cultures were then diluted 1:40 in agar supplemented with 5 mM $CaCl_2$, 5 mM $MgCl_2$ and 0.02/0.2 % (w/v) arabinose. Phage stocks were serially diluted eightfold in LB supplemented with 5 mM $CaCl_2$, 5 mM $MgCl_2$ and spotted on the culture-containing agar plates, which were incubated at 37°C overnight.

**JetABCD purification and reconstitution**

JetABCD proteins and complexes used in this study were produced in *E. coli* BL21. JetABC was co-expressed from a single vector, with JetA N-terminally tagged with 10His-TwinStrep-3C. JetD and the E248A mutant were expressed alone from a single expression vector with a C-terminal 3C-TwinStrep-10His-tag. A one liter culture of the strain containing the appropriate plasmid was grown in TB-medium at 37°C until the culture reached $OD_{600}$=0.5. For JetABC, the cultures were cooled down to 18°C and protein overexpression was induced by IPTG addition (0.4 mM final) for 16 hours. JetD(WT) and JetD(E248A) cultures were grown at 37°C until $OD_{600}$=0.5 and then induced for 20 hours by adding IPTG with final concentration of 0.08 mM at 16°C.

Cells were harvested by centrifugation and resuspended in lysis buffer (Tris pH 7.5 50 mM, NaCl 300 mM, glycerol 5 % (v/v), imidazole 25 mM) freshly supplemented with PMSF (1 mM) and β-mercaptoethanol (5 mM). Cells were lysed by sonication on ice with a VS70T tip using a SonoPuls unit (Bandelin), at 40 % output for 15 min with pulsing (1 sec on / 1 sec off). The lysate was then clarified by ultracentrifugation (40,000 g for 30 min). For JetABC, the supernatant was loaded onto a 5 mL StrepTrap column (Cytiva), followed by 5 column volumes (CV) washes using the lysis buffer. Protein was eluted with 4 CV of lysis buffer supplemented with 2.5 mM of desthiobiotin and 1,5 mL fractions were collected. Fractions containing the complex were then concentrated with Amicon Ultracentrifugal filter units (50 kDa cutoff; Millipore) and injected onto a Superose6 Increase 10/300 GL size-exclusion chromatography (SEC) column (equilibrated with 20 mM Tris–HCl pH 7.5, 250 mM NaCl and 1 mM TCEP). Output fractions were again concentrated to around 15-20 µM and flash frozen in small aliquots. For JetD/JetD(E248A), a 5 mL HisTrap column (Cytiva) was loaded with the lysate and washed by 10 CV of lysis buffer. Elution was performed with lysis buffer supplemented with 300 mM Imidazole and 3C protease was added to cleave the 10His-TwinStrep-3C-tag. The samples were dialyzed overnight at 4°C into dialysis buffer (20 mM Tris–HCl pH 7.5, 200 mM NaCl and 5 mM β-mercaptoethanol). JetD/JetD(E248A) was further diluted in Tris 20 mM pH 7.5, NaCl 100 mM buffer and loaded onto a 5 ml HiTrap Heparin column (Cytiva). After washing with 5 CV, bound material was eluted with 10 CV of Heparin elution buffer (20 mM Tris pH 7.5, 1000 mM NaCl and 5 mM β-mercaptoethanol). Fractions containing JetD/JetD(E248A) were then concentrated and further purified using a Superose 6 Increase 10/300 GL column (Cytiva) in buffer 20 mM Tris–HCl pH 7.5, 250 mM NaCl and 1 mM TCEP. Fractions containing JetD/JetD(E248A) were finally concentrated and flash frozen in liquid N2 for storage at -80°C .

Small scale reconstitution of JetABCD (for analytical purposes or cryo-EM) were performed by mixing 20 uL of JetABC and 20 µL of JetD from the frozen stock and incubated 10 minutes at 4°C. The mixture was injected on Superose6 Increase 3.2/300 with a SEC buffer containing either 20 mM Tris–HCl pH 7.5, 250 mM NaCl and 1 mM TCEP (for analytical purposes) or 10 mM Hepes-KOH pH 7.5, 150 mM KOAc, 2 mM MgCl2, and 1 mM TCEP (for cryo-EM). For biochemistry experiments, JetABCD was reconstituted by mixing 250 nM of JetABC dimer-of-dimers with 1000 nM of JetD in ATG buffer (10 mM Hepes-KOH pH 7.5, 150 mM KOAc, 5 mM MgCl2, and 1 mM TCEP) at room temperature for 5-10 minutes. ATG buffer was selected based on preliminary ATPase activity, ATP-dependent DNA binding and elution profile on analytical SEC. We note that DNA-cleavage competent JetABCD can also be reconstituted in a lower salt buffer (20 mM Tris–HCl pH 7.5, 30 mM NaCl, 2 mM $MgCl_2$).

**ATP hydrolysis assays**

The ATP hydrolysis activity of JetABCD was assessed by coupling to pyruvate kinase/lactate dehydrogenase at 37°C for 1 h in ATG buffer (lacking TCEP). 100 µl reactions were used, containing 1 mM NADH, 3 mM phosphoenol pyruvic acid, 100 U pyruvate kinase, 20 U lactate dehydrogenase, 1 mM ATP. The final concentration of JetABC/JetABCD (dimer-of-dimers) was 62.5 nM. JetABCD was reconstituted prior measurement by incubation of 625 nM of JetABC (dimer-of-dimers) with 1250 nM of JetD (dimer) in ATG buffer (lacking TCEP). 40 bp dsDNA was generated by annealing of complementary oligonucleotides and added to a final concentration of 125 nM. The reaction was monitored by measuring the absorbance evolution caused by NADH oxidation every minute at 340 nm using a Synergy Neo Hybrid Multi-Mode Microplate reader. The results were analyzed using GraphPad Prism V9.4.

**DNA cleavage experiments**

All reactions were performed in ATG buffer, supplemented with 1 mM ATP. Reconstituted JetABCD (WT or EQ, dimer-of-dimers, at 12.5 nM final conc. unless stated otherwise) was mixed with DNA substrate (pDonor, 8.5 nM or pJAZZ, 3.75 nM) in a final volume of 15 µL and incubated for 15 minutes at 37°C. After enzyme inactivation by addition of SDS (0.5 (w/v) % final) containing loading buffer and incubation for 10 minutes at 70°C, 15 µL of the reactions were loaded onto a EtBr-containing 1 % (w/v) agarose gel, ran at 5V/cm for 1 hour and bands visualized with on a transilluminator (U:GENIUS3 with a Syngene CAM-FLXCM-1 camera)).

pJAZZ was additionally pretreated with RecBCD to eliminate any linear DNA contaminations lacking hairpins that were routinely obtained from miniprep isolation. 1.5 µg pJAZZ DNA was pretreated with RecBCD (10 U, NEB) for 30 minutes at 37°C before enzyme inactivation for 30 minutes at 70°C.

For experiments involving restriction enzymes, 10 U were added per reaction in ATG buffer or appropriate reaction buffer. JetABCD titration experiments were performed with two-fold serially diluted (up to five times) JetABCD. For JetD titration, a two-fold serial dilution (up to six times) of JetD was prepared. For the linear DNA competition experiment, gDNA harvested from 1A700 *B. subtilis* was used. For each reaction, we added gDNA (max. 15x by mass compared to pDonor) which was two-fold serially diluted up to five times.

For experiments involving Proteinase K and RecBCD, 1 µg/ml and 10 U were used respectively. The RecBCD protection experiment was done by incubating a biotinylated PCR product (2.3 kb, 4.7 nM) pretreated with streptavidin-peroxidase conjugate (333 nM, Merck). For the re-circularisation experiments, reactions containing JetABCD/BsaI-treated plasmid was column purified (Epoch Life Sciences). 150 ng of the eluted DNA was then treated in T4 ligase buffer with T4 Ligase (5 U, Thermo Fisher) for 10 minutes at room temperature before heat inactivation for 10 minutes at 65°C.

Nicked pDonor DNA was prepared by mixing 10 µg of pDonor with Nt.BspQI (50 U, NEB). Reactions were incubated for 30 minutes at 50°C before heat inactivation for 20 minutes at 80°C. To generate relaxed pDonor, nicked pDonor was purified and further treated with T4 ligase (5 U).

**Cryo-EM**

<u>Sample preparation and data collection</u>

JetABCD complexes were freshly reconstituted in ATG buffer by size exclusion chromatography as described before (final concentration: about 0.5-0.7 mg/mL; 0.61-0.81 µM of dimer-of-dimers). Complexes were supplemented with ATP and β-octyl glucoside (final concentration of respectively 1 mM and 0.05 % (w/v)) and incubated 15 min at room temperature. Cryo-EM grids (Quantifoils R1.2/1.3

on 300 gold mesh) were freshly glow discharged in the EasyGlow device with 15 mA current. Then, 3 µL of JetABCD sample was applied on the grids mounted in a Vitrobot Mark IV. The Vitrobot was set to 10°C temperature in the chamber and 95 % humidity. Grids were blotted for seconds at blot force 5 without waiting and immediately vitrified in liquid ethane. Sample screening and an initial dataset acquisition was performed in a Glacios Cryo-TEM from Thermofisher Scientific (TFS). After few hours of data collection and validation of the sample/grid quality we moved the grids and collected high resolution data on a TFS 300kV Titan Krios G4i equipped with SelectriX energy filter at the DCI-Lausanne, UNIL. Images were collected on Falcon IV electron counting direct detection camera (TFS) in counted mode. The data were saved in the EER format using the EPU software (TFS) at a magnification of 165kx (pixel size is 0.726 Å at the specimen level) and a total dose of 40 electrons per square angstrom (e-/Å2) for each exposure, and defocus range of -0.8 to -1.6 micrometers. In total, 6404 movies were collected.

Data processing and model building

Initial data processing was performed on-the-fly using CryoSPARC-live with further refinement and classification using CryoSPARC (Punjani et al., 2017). In order to resolve high heterogeneity of the data the dataset was converted to Relion format with pyem scripts and further processed using RELION 4.0 (Scheres, 2012).

An overview of the data processing is shown in Figure S5. After motion correction with CryoSPARC-live, initial particle picking was performed using blob picker. Particles from good classes were used to feed the template picker to identify more particles. After several rounds of 2D classification, several *Ab initio* reconstructions (all giving very similar structures) were made using the particles from classes where the SMC shape of the complex was clearly visible (Figure S5). Homogenous refinement produced a map at an estimated global resolution of about 3 Å that was a starting point for initial stage of model building. We note that the coiled coils of the SMC on the side where poorly resolved. 3D variability analysis (Punjani and Fleet, 2021) indicated flexible motion in these regions of the map. Thus, the dataset was additionally treated in RELION. The first round of 3Drefinement was done using C2 symmetry. The following 3D classification without an alignment shows very high flexibility of the complex. In order to resolve the structure, we performed symmetry expansion. The refinement of symmetry expansion data improved the quality of the map and allowed us to build a model representing half of the dimer-of-dimers (Figure S5). In order to understand the nature of the flexibility we also perform 3D classification with alignment. The obtained 10 classes gives to us the idea how to complex may function. Finally, maps were combined to create a composite map that was used for final model building and refinement.

This map allowed the construction of a dimer-of-dimers model (containing two JetC dimers, a JetA dimer and two JetB dimers) by fitting two halves into the map. Note that we could not build a model for JetD nor for the JetC hinge domains and the adjacent coiled coils. To speed up modelling at the initial steps, we used structure predictions. AlphaFold was used to generate models for individual subunits or parts of JetABCD subunits, either alone or in complex (Evans et al., 2022; Jumper et al., 2021; Mirdita et al., 2022). Alphafold models were then rigid body fitted in the map using UCSF Chimera and ChimeraX (Pettersen et al., 2004; Pettersen et al., 2021) and rebuilt into the density using PHENIX real space refinement V1.20 (Afonine et al., 2018; Liebschner et al., 2019) and Coot V0.96 (Emsley et al., 2010). The model was validated using Molprobity (Davis et al., 2007).

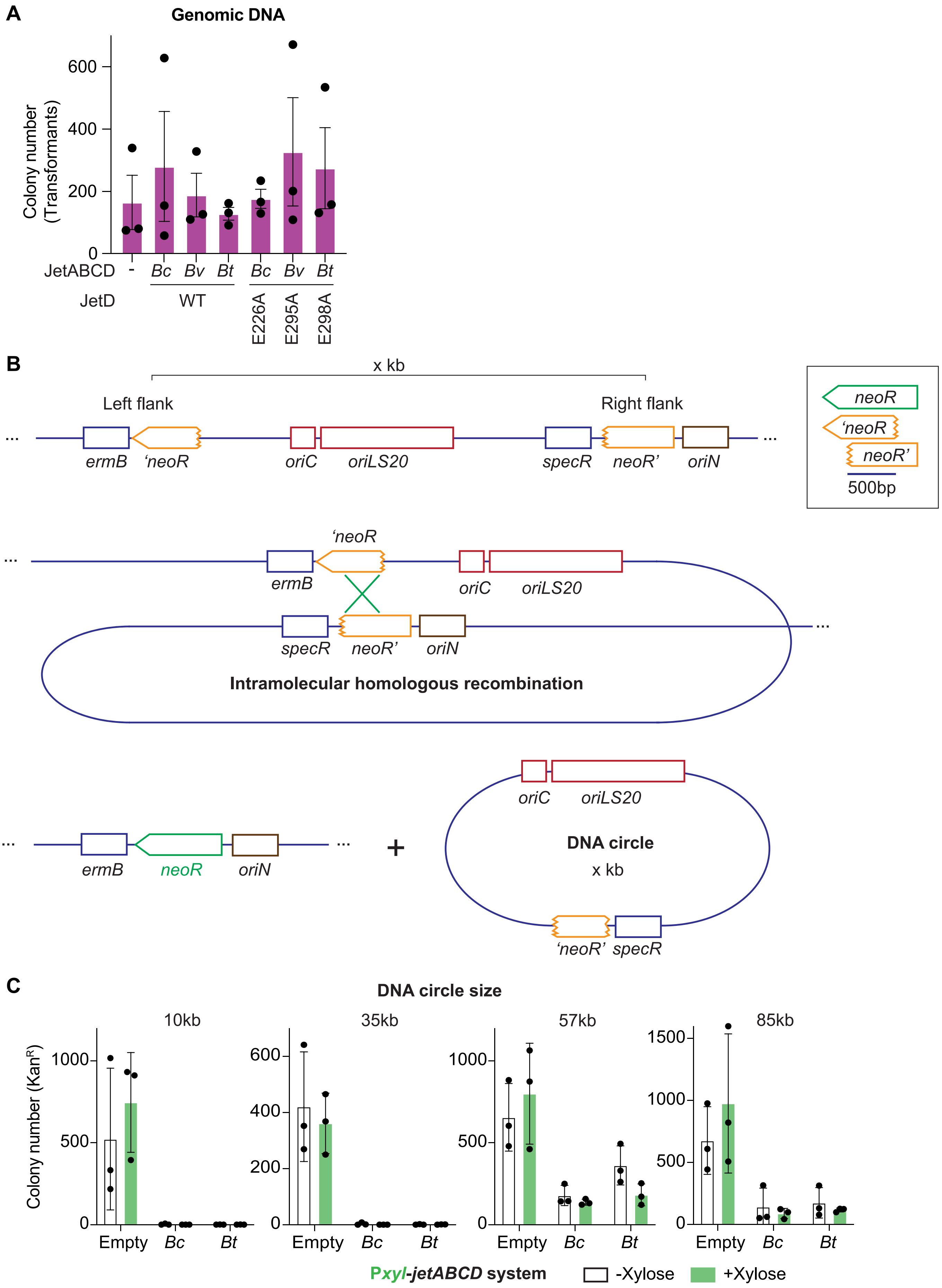
A
Genomic DNA
Colony number (Transformants)
600
400
200
0
JetABCD
-
Bc
Bv
Bt
Bc
Bv
Bt
JetD
WT
E226A
E295A
E298A
B
x kb
Left flank
Right flank
ermB
'neoR
oriC
oriLS20
specR
neoR'
oriN
neoR
'neoR
neoR'
500bp
Intramolecular homologous recombination
+
DNA circle
x kb
C
DNA circle size
10kb
35kb
57kb
85kb
Colony number (Kan^R)
Empty
Bc
Bt
Pxyl-jetABCD system
-Xylose
+Xylose

Figure S1

**Figure S1.** Related to Figure 1 and 2. **A)** Counts of antibiotic-resistant colonies for the indicated *B. subtilis* backgrounds after transformation with genomic DNA derived from BSG4618. Means and standard deviations from three independent experiments are shown. **B)** Detailed schematic of chromosome excision. Top panel: Depiction of flanks containing fragments of *neoR* (*neoR'* and *'neoR*) in direct orientation at defined distances from *oriC*. Box depicts the homology overlap of the *neoR* fragments. Middle: Depiction of intramolecular homologous recombination. Bottom: Depiction of the resulting excised DNA circle (right) containing the plasmid-derived *oriLS20*. (Left) the remaining chromosome DNA containing full-length functional *neoR* as well as exogenous *oriN*. **C)** Chromosome excision assays with *Pxyl-jetABCD*: Counts of kanamycin-resistant recombinants (containing DNA circles) indicated *B. subtilis* backgrounds with/without xylose (0.5% (w/v) final) addition. Means and standard deviations from three independent experiments are shown. Similar to results shown in Figure 1C except that JetABCD is cloned under a xylose-inducible promoter (*Pxyl*). We found that leaky expression from *Pxyl* (no induction) was sufficient to confer plasmid restriction and that promoter induction by addition of xylose did not alter the outcome substantially, implying that low protein levels are sufficient for robust activity and that higher levels are neither helpful nor obviously disruptive.

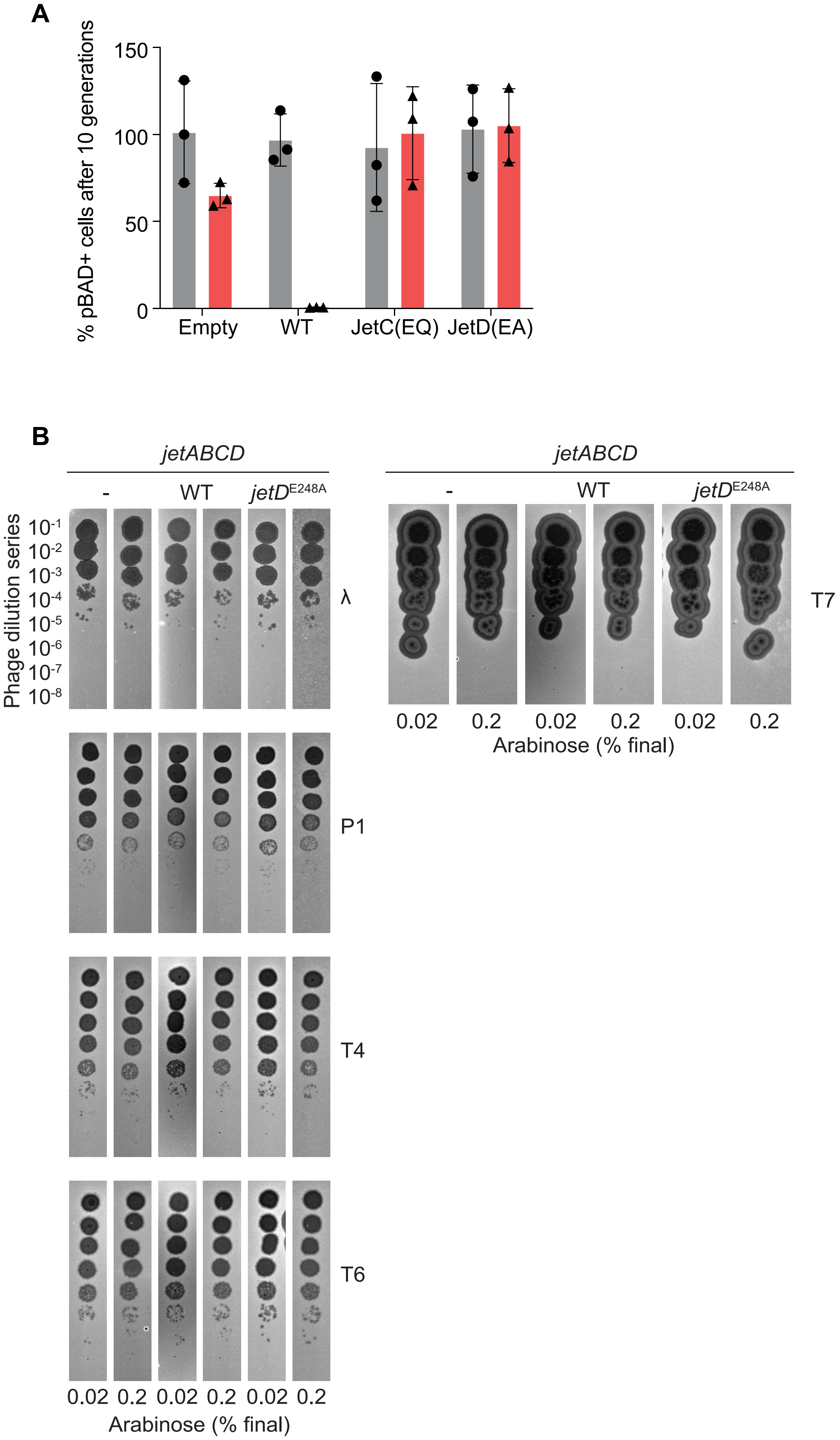


### Figure S2

**Figure S2.** Related to Figure 2. **A)** Graph showing the percentage of pBAD-containing *E. coli* cells in a cell population after ten generations without selection, with/without indicated mutant JetABCD induction by arabinose addition. Means and standard deviations from three independent experiments are shown. **B)** Phage protection assay for JetABCD-expressing *E. coli* strains. Plague formation from lawns of indicated *E. coli* strains upon incubation with indicated serially diluted phage.

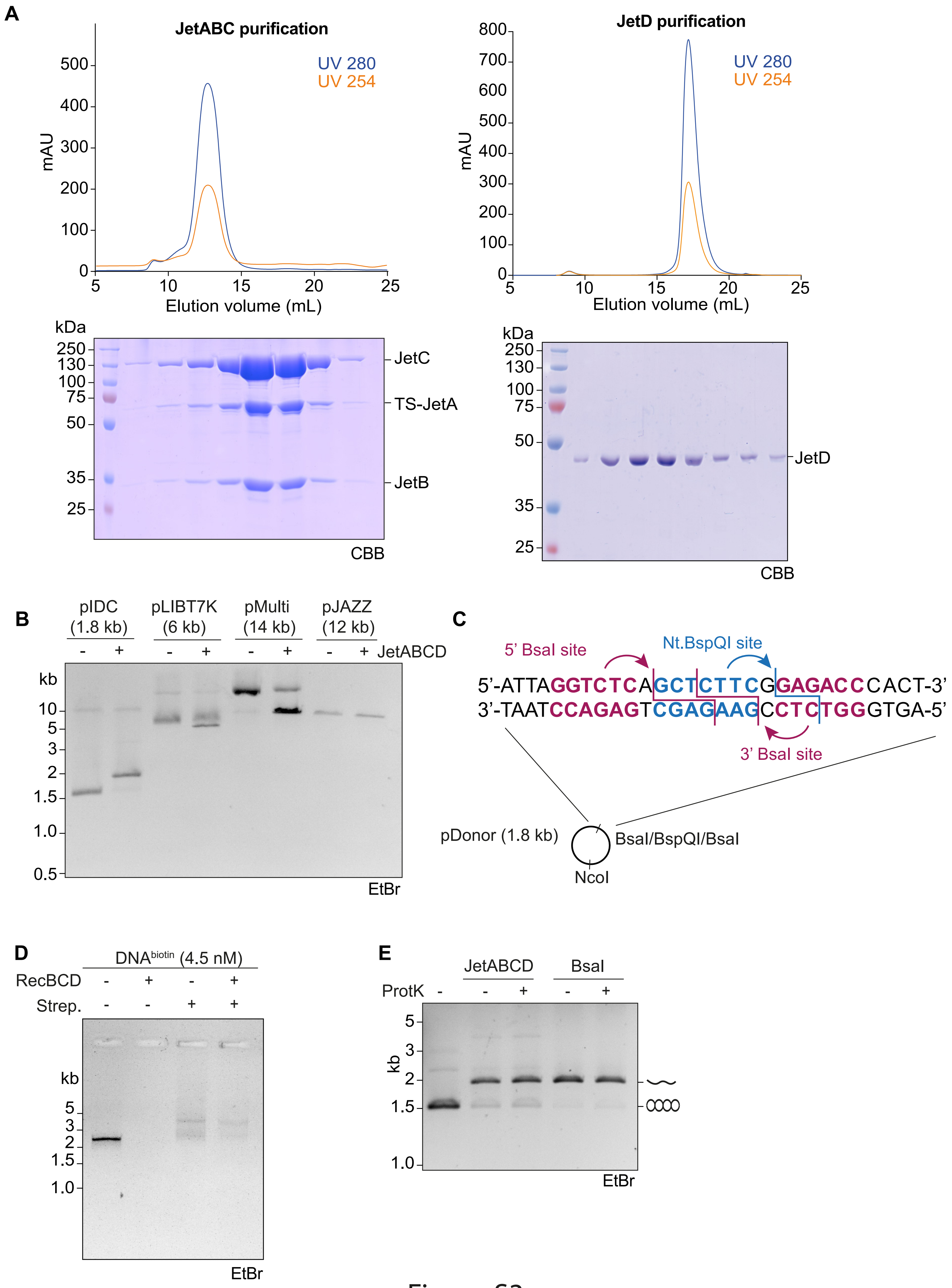


### Figure S3

**Figure S3.** Related to Figure 3. **A)** Purification of JetABC (left panel) and JetD (right panel). Top: elution profiles of the final gel filtration (Superose 6 Increase 10/300GL), bottom: peak fractions of the JetABC/JetD elutions were analysed by SDS-PAGE and Coomassie Brilliant Blue staining. **B)** Cleavage assay with reconstituted JetABCD and indicated plasmids. **C)** Schematic of pDonor showing its BsaI and BspQI sites. **D)** Gel migration of biotinylated DNA incubated with RecBCD and/or streptavidin. **E)** pDonor cleavage assay with JetABCD or BsaI-treated pDonor further incubated with or without proteinase K.

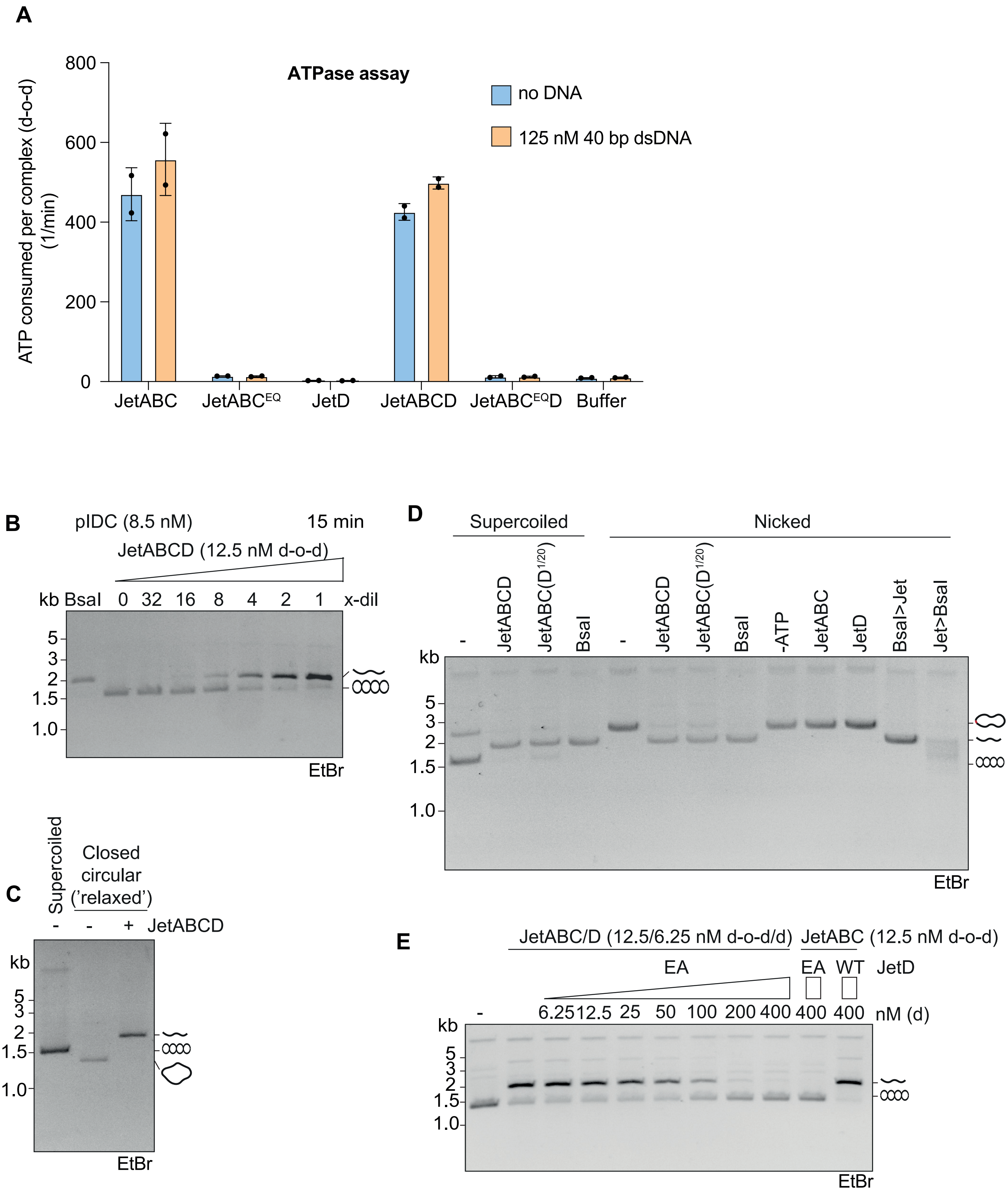


### Figure S4

**Figure S4.** Related to Figure 4. **A)** Enzyme-coupled ATP hydrolysis assay with JetABC (WT and EQ, dimer-of-dimers, 62.5 nM final conc.) and/or JetD (dimer, 125 nM final conc.). 40 bp dsDNA (at 125 nM final conc.) was added as indicated. **B)** DNA cleavage assay with decreasing amounts of JetABCD protein. As in Figure 4B with shorter incubation period (15 min). **C)** JetABCD cleavage assay of pDonor DNA in a relaxed conformation (after Nt.BspQI-nicking and subsequent T4 ligation). **D)** DNA cleavage assay with supercoiled and Nt.BspQI-nicked pDonor DNA at the indicated conditions. **E)** DNA cleavage assay with JetABC (12.5 nM of dimer-of-dimers) and JetD (6.25 nM dimer) and/or JetD(E248A) mutant at the indicated concentrations (max conc. 400 nM of dimer).

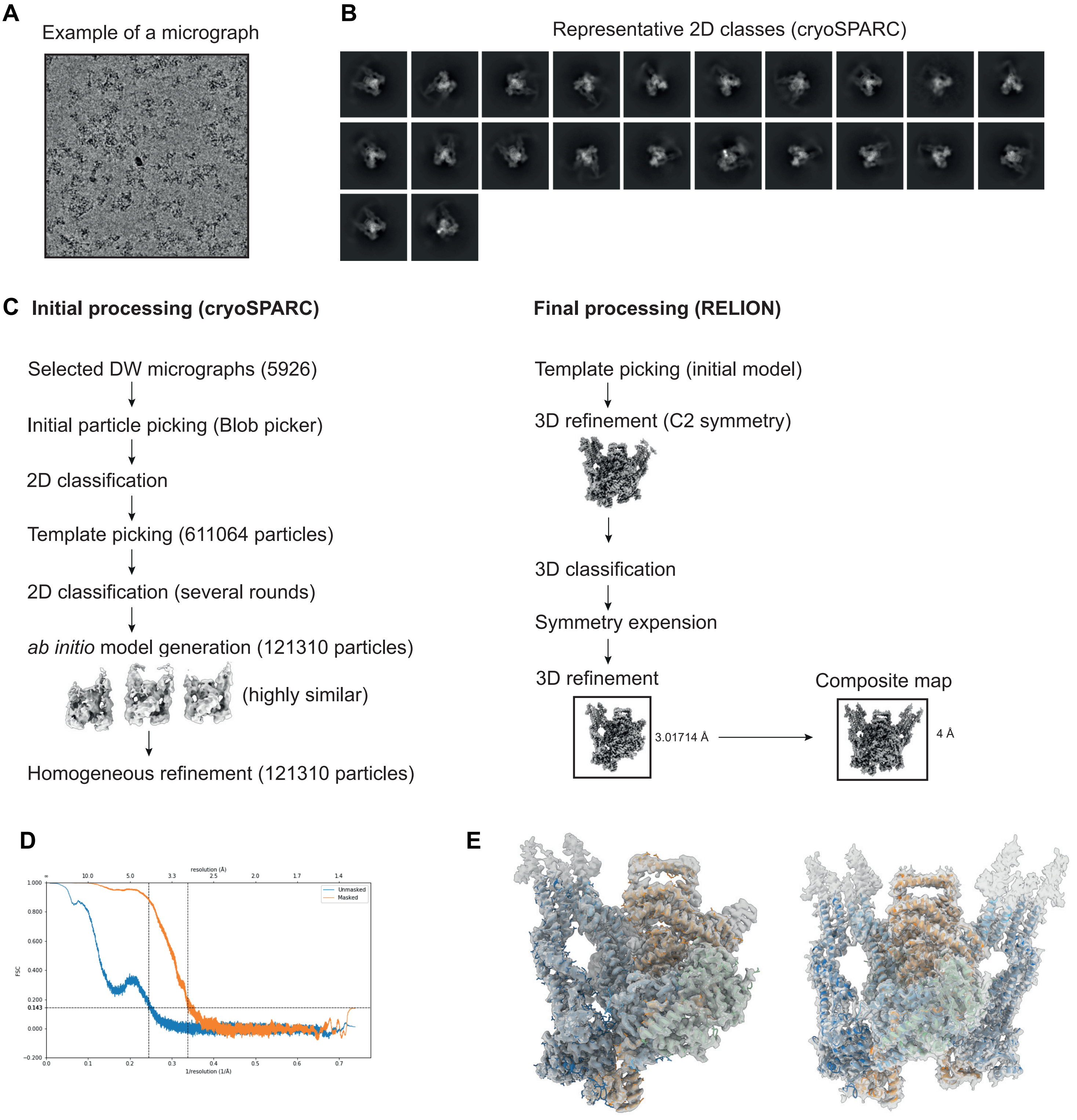


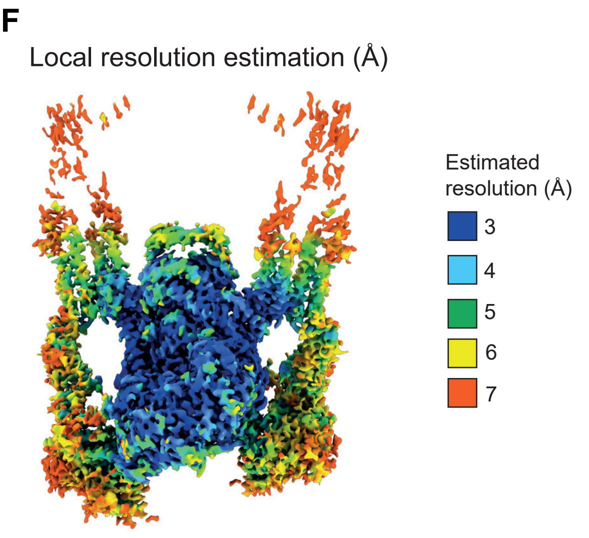


### Figure S5

**Figure S5.** Related to Figure 5. **A)** Example of a dose-weighted micrograph. **B)** Representative 2D classes from processing of the dataset with cryoSPARC. **C)** Cryo-EM data processing pipeline in cryoSPARC and RELION. **D)** Fourier shell correlation after 3D refinement in RELION (calculated using PHENIX). **E)** Models of JetABC placed in the maps (left: after RELION post-processing; right: in the composite map for the full dimer-of-dimers. **F)** Local resolution estimation of the map.

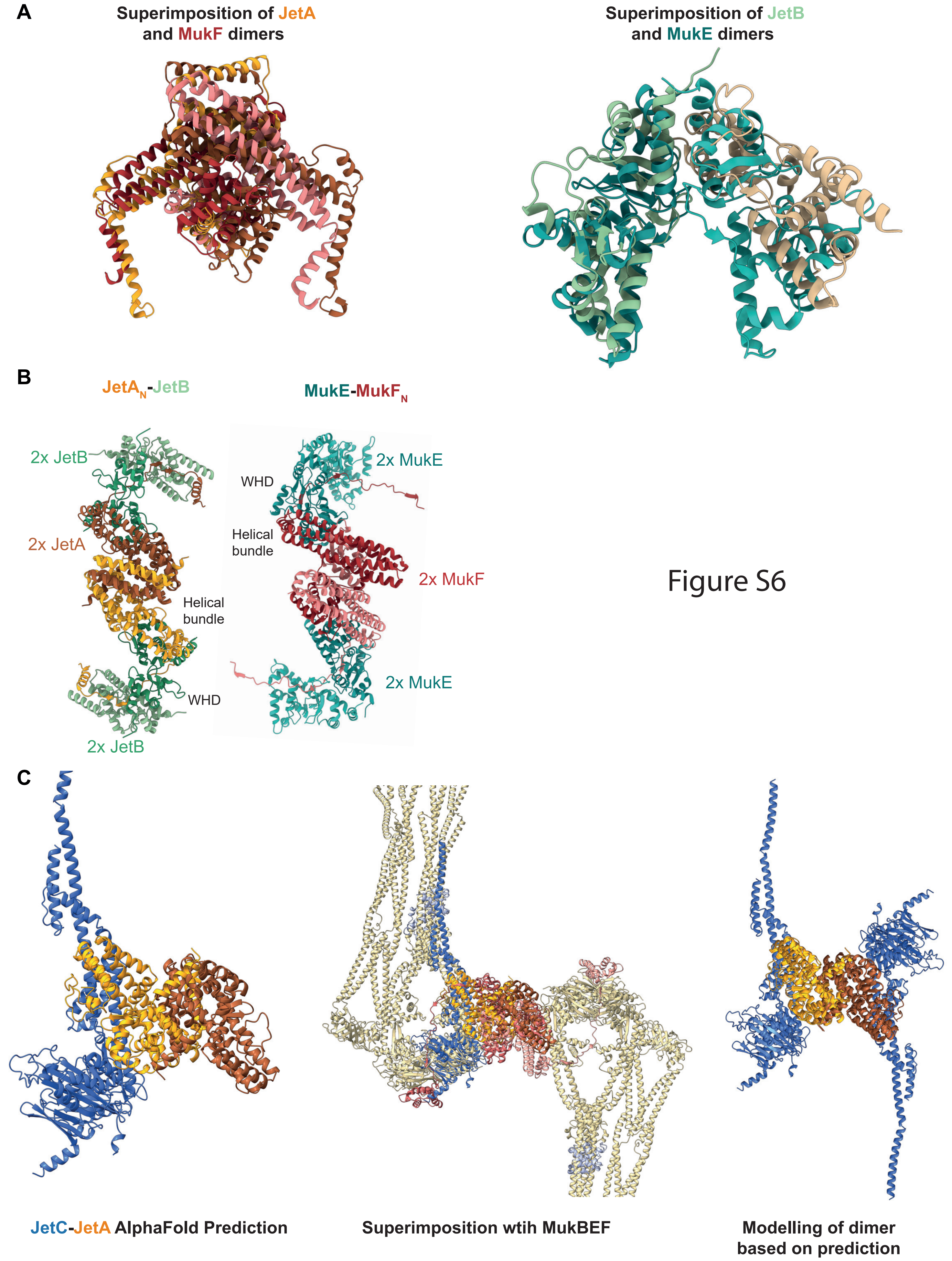


**Figure S6.** Related to Figure 5. **A)** Left: Superimposition of a JetAN dimer (cryo-EM model) and a MukFN dimer (PDB: 7nz4). Right: Superimposition of a JetB dimer (cryo-EM model) and a MukE dimer (PDB: 7nz4). **B)** Top views of JetB and MukE bound to JetA and MukF, respectively. Related to Figure 5B. **C)** Left panel: Alphafold prediction of a JetAN-JetC subcomplex. Middle panel: Superimposition of the JetA-JetC prediction with the MukBEF dimer-of-dimers (from PDB: 7nz4). Right panel: Modelling of a JetA-JetC dimer by self-superimposition of the AlphaFold prediction, revealing a hypothetical MukBEF-like open configuration.
